# Turmeric Phyto-NanoParticle (TPNP) enhances cellular bioavailability and anti-inflammatory effect of curcuminoids in human monocytes / macrophages

**DOI:** 10.1101/2025.07.21.665915

**Authors:** Bienvenue C. Habiyambere, D’Souza Kenneth, Victoria Northrup, Minji Kim, Victoria L. Nelson, Kyle R.D. Wells, Ghosh Anirban, Brunt R. Keith

## Abstract

The poor bioavailability of curcuminoids remains a major challenge to therapeutic use. This is largely due to their hydrophobicity, poor absorption, rapid metabolism, and short circulating half-life—limitations that are now being addressed through advances in nano- and micro-emulsion technologies. Curcuminoids and other water-insoluble phyto-polyphenols offer significant putative health benefits as anti-inflammatory, antioxidant, anticancer, radioprotective, and neuroprotective agents. Conventional emulsion-based delivery systems, such as liposomes, micelles, or solid lipid particles, rely on various emulsifying surfactants and/or excipients, some of which may themselves pose health risks. Here, we establish a novel class of all-natural, additive-free, oil-free, and emulsion-free Turmeric Phyto-NanoParticles (TPNPs) formulated directly from turmeric rhizomes and tested in a human monocyte/macrophage cell model to assess bioavailability kinetics and the efficacy of antioxidant and anti-inflammatory potential. TPNPs are enriched with curcuminoids (24.85% by mass), form a homogeneous nanoparticle distribution, exhibit higher antioxidant capacity, and demonstrate significantly improved cellular uptake in both monocytes and macrophages compared to conventionally purified curcuminoids. Favourable cellular pharmacodynamic anti-inflammatory effect of TPNPs was shown by increased levels of the cytoprotective enzyme heme oxygenase-1 (HMOX1), and a more effective reduction in lipopolysaccharide (LPS)-induced tumor necrosis factor (TNF) secretion compared to conventional curcuminoids. TPNPs could thus serve as a stable, non-synthetic, excipient-free formulation for safe and effective delivery of curcuminoids by nanocarriers for inflammatory conditions.

## 1. Introduction

The use of turmeric (*Curcuma longa*) as a medicinal natural product dates back approximately 6,000 years in the Indian subcontinent, from where its use spread across parts of Africa, China, and later Europe through Arab trade networks[1]. In modern contexts, turmeric and its derivatives are widely used across pharmaceuticals, nutraceuticals, food additives, dietary supplements, and cosmetic formulations[1]. Curcuminoids—a group of bioactive polyphenols derived from turmeric—hold significant challenges as active pharmaceutical ingredients (APIs) due to their poor bioavailability. This is primarily due to poor absorption, rapid metabolism, and a short circulating half-life. From antiquity to modern medical research, curcuminoids derived from the turmeric root have shown anti-inflammatory and anti-oxidative effects. In large part, this is attributable to its natural antioxidant properties and the commensurate upregulation of the cytoprotective enzyme heme oxygenase-1 (HMOX1), which is involved in pleiotropic metabolic effects and signalling[2].

When antioxidant defenses are overwhelmed, oxidative stress can broadly compromise cellular integrity, contributing to damage of membranes, proteins, and nucleic acids, and promoting downstream inflammatory and apoptotic signaling pathways. Human HMOX1 is a crucial enzyme in cellular stress response and plays a central role in immunological and inflammatory homeostasis, particularly in macrophages. It is featured in the transition state pro-inflammatory M1(-like) and anti-inflammatory M2(-like) macrophage phenotypes[3]. HMOX1 catalyzes the degradation of heme into biliverdin, carbon monoxide, and free iron—all of which possess cytoprotective and anti-inflammatory properties in various cell types, including monocytes/macrophages[3]. Biliverdin is subsequently converted into bilirubin, a potent antioxidant capable of mitigating oxidative stress by neutralizing reactive oxygen species (ROS). By promoting M2 polarization, HMOX1 supports inflammation resolution and tissue repair and is associated with improved cardiovascular function and reduced atherosclerotic risk[4]. TNF is a pleiotropic cytokine and a key orchestrator of inflammatory responses, produced primarily by cells of the monocytic lineage, such as macrophages, as well as other immune cells, including neutrophils and T cells [4]. Elevated TNF levels are implicated in rheumatoid arthritis, inflammatory bowel disease, psoriatic arthritis, and psoriasis, where they drive chronic inflammation and tissue pathology while concurrently impairing endothelial function and promoting vascular pathology[5]. Curcuminoids have been shown to induce HMOX1 expression[6][7], reduce TNF–driven inflammatory signaling in endothelial cells[8], and support anti-inflammatory M2 macrophage polarization[9], collectively contributing to immunomodulation and inflammation control. Curcuminoids exhibit cytoprotective effects via multiple molecular mechanisms: (i) scavenging of harmful free radicals[10]; (ii) induction of endogenous antioxidant enzymes and pathways, such as HMOX1[11]; and (iii) suppression of inflammatory cytokines, including TNF[12]. Curcuminoids have demonstrated beneficial effects across a range of disease contexts, including metabolic dysfunction, tissue repair, hepatic and cardiac toxicity, inflammatory bowel diseases, drug resistance, and immune modulation[13][14].

Despite extensive investigation, prior studies have inadequately addressed fundamental analytical limitations associated with curcumin, leading to misinterpretation of biological activity and poor clinical translatability. In particular, curcumin’s intrinsic fluorescence and absorbance properties, its tendency to spontaneously self-assemble into colloidal nanoparticles from reference standards, and its chemical instability have frequently confounded biological assays and sensitive methods used for active pharmaceutical ingredient (API) quantification. The absence of rigorous quality control, physicochemical characterization, and transparent disclosure of these confounders has further impeded the rational development of curcumin as a therapeutic agent[15]. Curcumin’s physicochemical properties—including poor aqueous solubility, instability, and intrinsic fluorescent activity—can produce assay interference that is often misattributed to biological effects[15], underscoring the necessity for systematic optimization and reporting of nanoparticle size and surface charge under varying biological media and storage conditions. Accordingly, accurate determination of curcuminoid API content in nanoparticle formulations requires validated analytical approaches, with reference-based quantification achievable by absorbance, fluorescence, or high-performance liquid chromatography (HPLC). The latter being the gold standard for reliable substrate profiling and translational rigour.

A wide array of curcuminoid encapsulation / formulation platforms—such as liposomes, micelles, and lipid-based nanoparticles—have been developed to enhance bioavailability, stability, and biocompatibility[16]. Though current nano- and microencapsulation systems rely on specialized carrier oils, emulsifiers and surfactants, many pose toxicity risks. To prepare water compatible (micro and nano) formulations of hydrophobic compounds/drugs, either (i) surfactants are mixed with hydrophobic compounds to form a hydrophobic core and water compatible outer layer, or (ii) hydrophobic compounds dissolved in carrier oil form a core structure surrounded by water compatible layer of emulsifiers, also known as oil-in-water formulations[17][18][19]. Carrier oils are usually sourced from plants and are challenging for oil-in-water formulations due to their high viscosity, variable composition, processing difficulties, limited drug-loading capacity, and poor shelf stability [19]. Emulsifiers are a subclass of surfactants used to stabilize oil-in-water formulations. Increasing evidence links dietary emulsifiers and surfactants—used as stabilizers or delivery agents—to adverse health and environmental effects, particularly with prolonged, low-dose exposure. These agents have also been shown to alter gut microbiota composition[20][21], contribute to metabolic disorders—including cardiovascular disease, hypertension, and type 2 diabetes[22][23][24] —and impair gut epithelial integrity or dysbiosis[25][26]. Chronic exposure to emulsifiers can be associated with long-term health burdens [27], and animal studies further suggest that certain emulsifiers can alter gene expression in brain cells, thereby influencing stress responses and blood pressure regulation [28][29]. Chemically, surfactants and emulsifiers are amphiphilic molecules that act as surface-active agents having a specific critical micelle concentration to form a micellar structure in water with a hydrophobic core[17][18][19]. Widely used non-ionic surfactants like Polysorbates (Tweens) have been reported to cause cytotoxic and genotoxic effects *in vitro*[27][28][29][30]. Cationic surfactants—such as cetyltrimethylammonium bromide, dimethyl-dioctadecylammonium bromide, and cetylpyridinium chloride—are essential in nucleic acid delivery systems but are also associated with significant cytotoxic potential[31][32][33]. Polyethylene glycol (PEG), is a commonly used polymer surfactant, and broadly approved for use in food, cosmetics, and pharmaceuticals due to its biocompatibility and favorable pharmacokinetic properties[34]. Nonetheless, frequent exposure to PEG-containing products elicited anti-PEG antibody responses in the general population [35], raising concerns about immunogenicity from cumulative exposure through food, drugs, and vaccines. In recent decades, an increasing number of clinical trials have examined multiple orally delivered curcuminoids for disease treatment, with promising results across a range of conditions [36][37][38][39]. Approximately half of these trials have focused on diseases involving chronic inflammation, including metabolic and musculoskeletal disorders, and have generally reported favourable therapeutic outcomes such as reduced levels of pro-inflammatory cytokines TNF and IL-6. Collectively, this highlights the desire for a reliable, reproducible, quality-assured, referenced nanocurcuminoid delivery platform that could improve bioavailability while eliminating reliance on carrier oils, emulsifiers, and surfactants, thereby reducing excipient-associated toxicity and immunogenic risk without compromising therapeutic efficacy.

In pharmacokinetics, bioavailability refers to the proportion of an administered dose of an active API that enters the systemic circulation, as well as the rate at which this occurs. The kinetics of absorption, metabolism, and elimination are key factors that influence effective bioavailability and the maintenance of steady-state plasma concentrations. Cellular pharmacodynamics refers to the assessment of the biochemical and physiological effects exerted by the API on living cells. It encompasses the characterization of cellular responses, including receptor interactions, signal transduction, and downstream functional outcomes that reflect the mechanism of action and therapeutic potential of these agents. Two principal approaches can be used to demonstrate the enhanced therapeutic efficacy of a colloidal nanoformulation relative to a reference standard: (1) achieving higher therapeutic outcomes using equimolar concentrations, or (2) attaining equivalent efficacy or higher at a reduced dose of the colloidal preparation. Preclinical studies frequently utilize *in vitro* models to assess cellular bioavailability and pharmacodynamics to predict and validate the therapeutic efficacy of APIs. Assessing the colloidal API in isolation using simplified chemical assays or *in vitro* models is inadequate for predicting therapeutic efficacy; evaluating both the API and its mechanism of action—here evidenced by HMOX1 induction and TNF suppression—is essential.

In this study, we evaluate the intracellular bioavailability and anti-inflammatory potential of a novel, all-natural Turmeric Phyto-NanoParticle (TPNP) nanoformulation that is free from emulsifiers, surfactants, and carrier oils. Importantly, the entire composition of TPNP is derived from the whole turmeric rhizome, ensuring a bio-native micelle matrix devoid of synthetic additives or delivery agents.

## 2. Materials and methods

### 2.1 THP-1 cell culture

THP-1 human monocytes were expanded from frozen stocks originally purchased from the American Type Tissue Culture Collection (ATCC) using media containing RPMI-1640 (ThermoFisher Scientific, Cat# A1049101) supplemented with 10% FBS (ThermoFisher Scientific, Cat# 12483020). Cells were maintained at 37°C in a 95% air, 5% CO_2_ humidified atmosphere and the media was changed every 3-4 days. THP-1 monocytes (suspension cells) were differentiated into adherent macrophages using a final concentration of 10nM phorbol 12-myristate 13-acetate (PMA) (VWR, Cat# 10187-494) in RPMI1640-10% FBS media for 16 h. After differentiation with PMA, the media was replenished with PMA-free RPMI-1640-10% FBS media and incubated for 30 h. Then the media was replenished with RPMI-1640-0.5% FBS media and incubated for 16h. Experiments were carried forward using RPMI-1640 with 0.5% FBS.

### 2.2 TPNP and Curcuminoid-standard Spontaneously Aggregating Particles (CSAP)

TPNP were manufactured and supplied by Pividl Bioscience Inc. (Canada) through a proprietary process as a sterile aqueous suspension in 0.1x PBS (pH 7.4) with 15% ethanol. TPNP are manufactured as bio-native micelle-type structures enriched with curcuminoids directly from turmeric rhizomes using a hydroethanolic process. The standard curcumin used was purchased from Sigma-Aldrich (CAT #C1386). **CSAP formation in cell treatment media:** The cell culture media used for treatment was RPMI-1640 supplemented with 0.5% FBS. The media was spin-filtered (300 kDa filters) to remove particles like extracellular vesicles and protein aggregates present in FBS. Curcumin powder (Sigma-Aldrich Cat#1386) was dissolved in 95% ethanol to make a stock solution (1.5 mM), which was used to prepare 50 µM curcuminoids equivalent CSAP in 300 kDa-filtered RPMI-1640-0.5% FBS media following incubation at 37 °C for 30 min with intermittent mixing; CSAP colloidal formation was observed. CSAP formation in aqueous media of RPMI-1640 and 0.1x PBS, with or without 0.5% FBS were evaluated.

### 2.3 Nanoparticle characterization

Nanoparticle properties were determined using Dynamic Light Scattering (DLS) and Tunable Resistive Pulse Sensing (TRPS) using a Zetasizer Nano ZSP (Malvern) and qNano Gold (Izon) instruments, respectively. TPNP were diluted 10 or 15x (in 0.1x PBS) and 10,000x (in deionized water) for DLS and TRPS, respectively.

### 2.4 Total curcuminoid quantitation using fluorescent spectroscopy and HPLC equipped with an optical diode array detector (DAD)

Synergy H4 Hybrid Reader (Biotek) was used for fluorescent spectroscopy. Curcumin powder was measured and dissolved in 80% ethanol to make two stocks of 1.5 mM curcuminoids equivalent and serially diluted to 1000, 100, 10 and 1 nM in 80% ethanol to produce a standard curve with excitation at 420 nm and emission capture at 570 nm. TPNP samples were diluted 3125x in 80% ethanol for the above quantitation. The fluorescent spectroscopy method was further validated with HPLC-DAD. To quantify if three curcuminoids are present in standard curcuminoid powder and TPNP formulation using HPLC-DAD, a ternary isocratic chromatographic method, coupled with optical diode array detection monitoring compound-specific wavelengths was used. Stock solutions of the three individual curcuminoid standards (Curcumin, Desmethoxycurcumin and Bisdemethoxycurcumin) at 1000 μg/mL were prepared in acetone, followed by dilution to 100 μg/mL (100 ppm) in acetonitrile and used to perform HPLC-DAD method optimization and to acquire spectra for the individual compounds. Further validation was performed by mixing three individual curcuminoid standards at 100 μg/mL and used for daily instrument calibration. The three reference standards obtained for this work had a purity equal to or greater than 98%. A final ternary isocratic method consisting of 30% acetonitrile, 25% methanol and 45% water, supplemented with 0.1% phosphoric acid was employed to separate the three curcuminoids sufficiently and minimize secondary interaction of the curcuminoids with the nanocarrier. DAD scanning range was set to 210–800 nm with 1 nm increments, and data acquisition was performed using MassLynx 4.2 with TargetLynx software.

### 2.5 Transmission electron microscopy (TEM)

5 or 10 µL TPNP was directly deposited onto 200 mesh formvar/carbon grids and allowed to incubate at room temperature for 10 min before removing the excess liquid by wicking with filter paper. The grid was then sequentially washed twice for 2 min by floating sample-side down on two separate 100 µl of ddH_2_O (0.22 µm filtered) droplets on parafilm. Contrast staining was done by floating the sample-side down on 1% uranyl acetate (Fisher Scientific Company) droplet for 1 min. Grids were imaged with a STEM3+ detector on a Scios 2 DualBeam (Thermofisher Scientific) at 80,000x magnification and 30 kV accelerating voltage.

### 2.6 Antioxidant capacity

The total antioxidant capacity was determined according to the recommended procedures of the manufacturer of the Ferric Reduction Antioxidant Potential (FRAP) assay kit (ThermoFisher, Cat# EIAFECL2). In brief, 10 mM Ferrous chloride was diluted in 1x acetate buffer to prepare Fe^2+^ standards of 0, 31.25, 62.5, 125, 250, 500 and 1000 µM by serial dilution. All samples were diluted in 1x assay buffer. Fresh stock of 62.5 µM of each of Curcuminoids Standard, N-Acetyl-L-Cysteine [NAC] (Amresco, Cat# 0108-25G) and Gallic Acid (GA) were made in 1x assay buffer as references. A fresh Fe^2+^ standard curve was prepared for each assay. The absorbance was read at 560nm using a Synergy H4 Hybrid Reader (Biotek).

### 2.7 Untargeted Metabolomic Analysis Using Nano-LC-MS/MS

TPNP was solubilized in 99% ethanol (1:10) and was separated on a 150 mm x 75 μm Easy SprayTM C18 column (Thermofisher Scientific, Cat# ES900) using a gradient elution with water and acetonitrile. Samples were loaded onto the column for 60 min at a flow rate of 230 nL/min. Metabolites were separated using a gradient that starts with 100% water and 0% acetonitrile in the presence of 0.1% FA and were separated for 35 min with a final gradient to 100% acetonitrile. Separation continued for an additional 10 min using 100% acetonitrile, followed by a 5 min run, then gradually decreasing to 0% acetonitrile and 100% water and maintained that ratio for 10 min at the end. Eluted metabolites were directly sprayed into a mass spectrometer using positive electrospray ionization (ESI) at an ion source temperature of 250℃ and an ion spray voltage of 2.1 kV. The full-scan MS spectra (m/z 80-1200) were acquired at a resolution of 70,000. Precursor ions were filtered according to the monoisotopic precursor selection, with charge state of 1 and 2, and dynamic exclusion of 10s. The automatic gain control settings were 3×10^6^ for full FTMS scans and 1×10^5^ for MS/MS scans. Precursor ions were isolated using a 2 m/z isolation window and fragmented with a normalized high energy collision of 35%.

Spectral Data Preprocessing: Each data file was uploaded onto the opensource MzMine3 software for preprocessing. Masses for each sample were detected using a 5.0E5 noise threshold limit for both MS1 and MS2 spectra. ADAP Chromatogram Builder, a module of MzMine3, was then used to peak pick based on the following settings: minimum highest intensity 5.0E5 and noise of 5.0E5 with 5 data points per peak. Once the chromatograms were created, the Local Minimum Resolver module was then used to clean up the features and link the MS1 and MS2 spectra using “MS/MS scan pairing”. The chromatogram threshold was set to 90% with a minimum search range RT of 0.03 and a minimum relative height of 50%. The ratio was 1.2 and the peak duration was set to 1 minute. Deisotoping was then performed where the retention time tolerance was set to 0.5 min and the m/z tolerance was set to 5 ppm. The peaks that were in the blank were also subtracted from the sample using the default setting for data points and a fold increase of 150%, meaning that true peaks were identified if the area was 150% more than the noise level.

Metabolite annotation was conducted using the opensource Sirius 4 CSI: Fingerid software. The .mgf file of spectral data, exported from MzMine3, was imported into Sirius 4. Default parameters were used for identification, using “Orbitrap” as the type of data and the following database were used: BioDataBase, Biocys, KNApSAcK, Natural products, Plantcys, YMDB and GNPS.

### 2.8 Immunoblotting and Confocal microscopy

For immunoblotting, THP-1 were seeded at a density of 1-1.5 x 10^6^ cells on 35 mm dishes and differentiated into macrophages. Cells were then treated with CSAP or TPNP for comparative analysis of protein targets. Protein lysate samples and ladder were separated via SDS-polyacrylamide gel electrophoresis (Bio-Rad Cat# 5671095) and transferred onto nitrocellulose membrane (0.2µm, Bio-Rad# 1620112).

Membranes were reversibly stained with MemCode (ThermoFisher Scientific Cat#24580) before immunoblotting. 5% milk was used to block for 45 min and incubated with HMOX1 primary antibody (CST Cat#5853, 1:1000 dilution in 1% skim milk in TBST with 1% sodium azide overnight). Membranes were washed in 1x TBS-T, followed by incubation with Goat Anti-Rabbit Horseradish Peroxidase-Conjugated Secondary Antibody in 5% milk (1:2000) for 2 h. The signal was detected by enhanced chemiluminescence (ThermoFisher Scientific #34076) and digital imaging. Western blot analysis was performed using Image Lab software and values were obtained by measuring the protein target relative to the total protein (MemCode). For confocal microscopy, THP-1 cells were seeded at a density of 1×10^5^ cells/well on sterilized coverslips placed in 6-well plates and differentiated to macrophages. Cells were then stained with a 1µL:1000µL dilution of Phalloidin 647 (VWR, Cat# 89427-130) for 30 min. Hoechst 33258 (Thermofisher Scientific, Cat#62249) was used at a 0.5µL:10 000µL dilution to stain nucleus for 1 min and finally mounted/sealed on a microscope slide using PermaFluor mounting media (Thermofisher Scientific, Cat# TA030FM). Confocal images were acquired using Zeiss LSM 900 Axio Observer (equipped with Airyscan 2).

### 2.9 Flow cytometry

We used curcuminoids’ autofluorescence to measure comparative curcuminoid accumulation in treated cells. THP-1 cells were treated as indicated in the result sections followed by fixing in 2% formalin in FACS buffer (0.1% BSA in 1x PBS) and subsequently washed twice with 1x PBS. Cells were then gated and stored (75,000 cells) using the GalliosTM 10-color Flow Cytometer (Beckman Coulter) and curcumin fluorescence was assessed in the FL2+ channel.

### 2.10 Cell viability assay

Cell viability assay was performed as per supplier recommendation of PrestoBlue^TM^ Cell Viability Reagent (Invitrogen, Cat#A13261). Spectral fluorescence was measured using a Synergy H4 Hybrid Reader (Biotek) with excitation and emission wavelengths of 560 and 590 nm respectively. Cell viability was calculated by subtracting the fluorescence at 560/590nm media-only control or CSAP/TPNP (i.e., to correct for curcumin’s autofluorescence between 420nm-600nm) from the respective cell treatment groups and normalizing values to the mean viability of the untreated controls.

### 2.11 ELISA

For TNF measurement in the cell culture media, the ELISA MAX^TM^ Standard-set kit (BioLegend, Cat#CA430201-BL) was used according to the manufacturer’s protocol and absorbances were read at 450 nm using a Synergy H4 Hybrid Reader (Biotek).

## 3. Results

### 3.1. Turmeric Phyto-NanoParticle (TPNP) characterization

Each batch of TPNP was stored at 4-8°C for three months and batch-to-batch hydrodynamic diameter (nm) and polydispersity index (PDI) was verified after 10x dilutions in 0.1x PBS (Table 1) using DLS. To assess long-term shelf-stability, the above TPNPs were dried using vacuum assisted freeze-drying (−40°C) with sucrose (10% w/v, a cryo-drying excipient) to a powder form (Figure 1A) and immediately sealed in airtight containers, stored at room temperature in dark and dry environment. For rehydration, 0.1g of the dried TPNPs was dissolved in 1 mL of ddH_2_O, followed by a further 10x dilution in ddH_2_O. The rehydrated TPNP powder showed an increased size compared to undried/hydrated TPNP but still maintained homogeneity for up to 24 months, as shown by low PDI values at 6, 9 and 24 months (Table-1). To assess nanoparticle-induced turbidity, a red laser was directed through rehydrated freeze-dried TPNPs in water (left beaker) and water alone (right beaker) (Figure-1B). TPNPs morphology was visualized by TEM and found to be spherically shaped (Figure-1C). The average hydrodynamic diameter distribution of TPNPs shown in Figure-1D was measured by DLS after 10x dilution in 0.1x PBS and a single intensity peak (177 ± 0.964 nm) was observed, demonstrating particle size homogeneity (PDI= 0.111). The uniformity of particle population was also shown by the correlation coefficient curve (Figure-1D inset).

**Figure 1:**
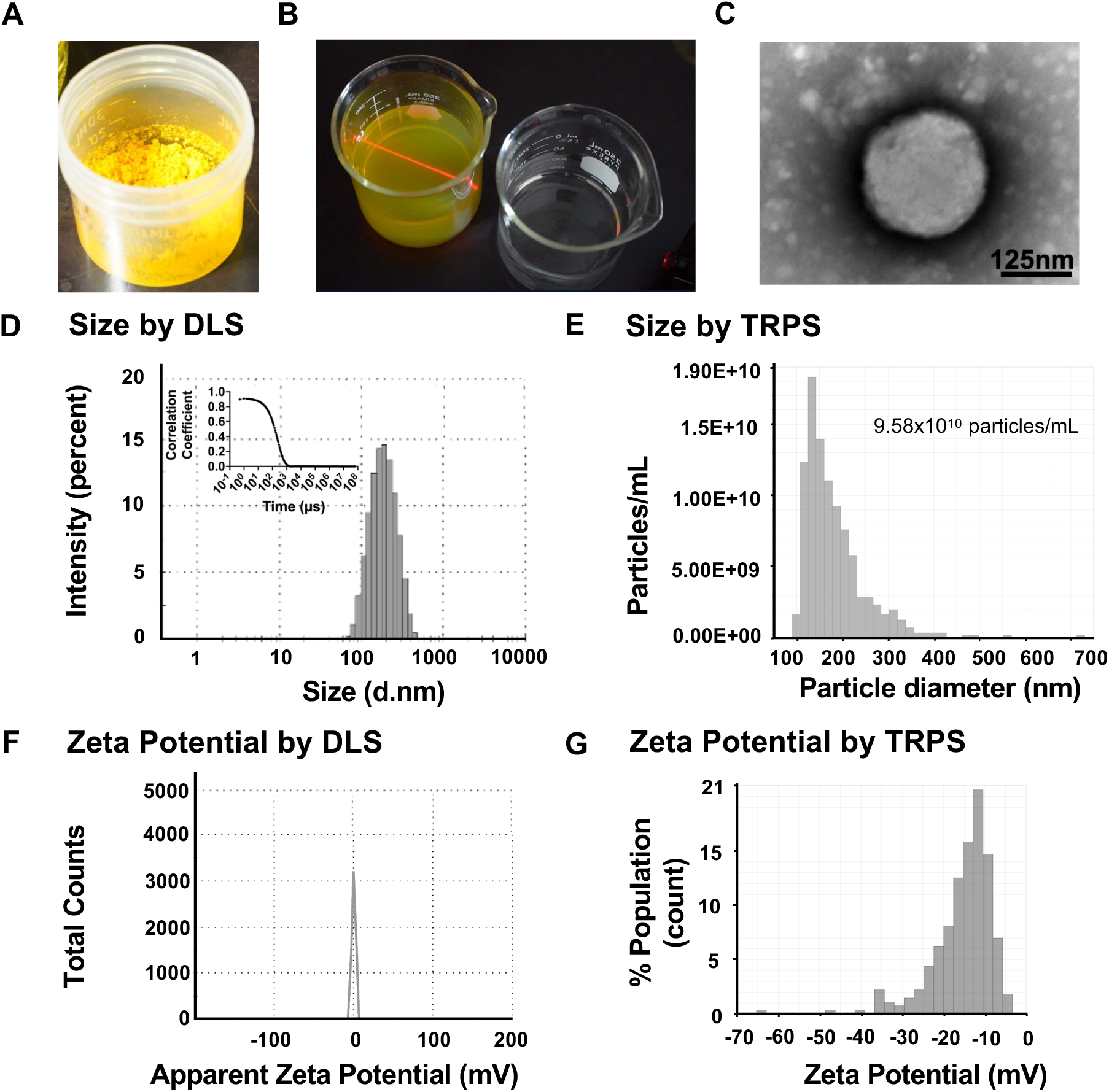
Biophysical characterization of TPNPs confirms nanoparticle uniformity and conformity, with a near-neutral negative surface charge. (A) Freeze-dried TPNP stored at room temperature. (B) Rehydration of freeze-dried TPNP in water (left beaker) compared to water alone (right beaker), with a red laser beam passed through both beakers to demonstrate colloidal turbidity. (C) A representative image of TPNP using Transmission Electron Microscopy showing slightly smaller size to DLS/TRPS readings likely due to dehydration protocol required for TEM imaging. (D) Particle size distribution of TPNP using DLS where the inset represents the particles monodisperse correlation coefficient curve (a measure of decaying exponential function), where x-axis values are represented on log10 scale. (E) Particle size distribution of TPNP using TRPS, (F) zeta potential of TPNP using DLS and (G) zeta potential of TPNP using TRPS.

**Table-1:** TPNPs exhibit stability and maintain particle homogeneity following long-term storage. The table shows average hydrodynamic size (d.nm) and PDI values obtained by DLS for two different TPNP batches (undried/hydrated and dried/rehydrated). The freeze-dried and rehydrated batch was monitored over a 24-month period.

| Formulation type | Time | Average hydrodynamic size (d.nm) | PDI |
| --- | --- | --- | --- |
| Hydrated TPNP | Batch #1 | 173 ± 0.762 | 0.131 |
|  | Batch #2 | 177 ± 0.964 | 0.111 |
| Freeze-Dried TPNP (Batch #1) | 6 months | 214 ± 2.86 | 0.055 |
|  | 9 months | 235 ± 1.76 | 0.128 |
|  | 24 months | 223 ± 3.76 | 0.254 |

TPNPs was also analysed using Tunable Resistive Pulse Sensing (TRPS)-based particle analysis. Figure-1E shows TRPS of TPNPs diluted 10,000x in deionized water and found to contain about 9.58 x 10^10^ particles / mL with a mean particle diameter 176 ± 67.9 nm and a mode particle diameter of 132 nm. Using DLS, TPNP showed a net zeta potential of −0.189 mV when diluted in an ionic environment (10x dilution in 0.1x PBS) (Figure-1F).

Zeta potential of TPNPs was measured using TRPS following a 10 000x dilution in deionized water, yielding a mean value of −15.0 mV and a mode of −11.7 mV (Figure-1G). The discrepancies of zeta potential value of DLS data in comparison to TRPS data are likely due to the presence of sufficient buffering salt from 0.1x PBS (pH7.4) used as diluent in DLS, which neutralized charges, whereas 10,000X dilution in deionized water did not have sufficient buffering salt.

### 3.2 Curcuminoids Standard Spontaneous Aggregating Particle (CSAP) formation in cell treatment media

Prior studies have failed to account for curcuminoids’ inherent properties, such as their spontaneous formation of nanoparticles in cell culture media[40], which creates inaccurate interpretations and confounds clinical translation. We validated that curcumin–used here as a pharmacological reference standard–would spontaneously form particles by aggregation in cell treatment media. The hydrodynamic diameter of the CSAP formed in the cell treatment media (RPMI-1640 media + 0.5% FBS) exhibited an average size of 134.2 ± 0.9 nm and PDI of 0.146 by DLS (Figure S1A). To determine whether curcuminoids could form similar-sized aggregating particles, we compared curcuminoids in 0.1x PBS or RPMI-1640 media in the absence or presence of FBS. In RPMI-1640 without FBS, curcuminoids formed larger aggregates, averaging 410.1 ± 8.7 nm with a PDI of 0.260 (Figure S1B). In contrast, when diluted in 0.1x PBS with 0.5% FBS, the average hydrodynamic diameter was substantially smaller at 83.6 ± 1.2 nm and with a lower PDI of 0.198 (Figure S1C). In 0.1x PBS alone, curcuminoid aggregation resulted in particles averaging 1088 ± 54.0 nm with a PDI of 0.244 (Figure S1D). All CSAP aggregates in the above examples of treatment media either precipitate or stick to the container wall after 30 h of incubation at room temperature (data not shown). These results indicate that the presence of complex biological fluids containing proteins, lipids and metabolites stabilizes the curcuminoid colloidal particle, resulting in smaller particle sizes in aqueous environments.

### 3.3 TPNP in cell treatment media

To evaluate whether TPNPs undergo changes in particle size in cell-treatment media (RPMI-1640 + 0.5% FBS) compared to their original size of 177nm and PDI of 0.111 in 0.1x PBS, TPNPs were diluted to make 50 µM curcuminoids-equivalent (matching the concentration used in the CSAP study) in media with or without FBS. In RPMI-1640 + 0.5% FBS, the average hydrodynamic diameter increased to 223.8 nm ± 3.2 nm with a PDI of 0.121 (Figure S1E), whereas TPNP diluted in RPMI-1640 without FBS exhibited a smaller diameter of 171.1 ± 0.5 nm and PDI of 0.04 (Figure S1F). These results suggest that complex biological molecules present in FBS may adsorb onto the nanoparticle surface, forming a biological corona that increases the apparent hydrodynamic size.

### 3.4 Pharmaceutical equivalent curcuminoids in TPNP

For a rapid and inexpensive 96-well assay, CSAP and TPNP were prepared in 0.1x PBS and scanned across the UV-visible spectrum (250 nm-800 nm) at 1 nm intervals. The absorbance maximum was around 430 nm for both samples (Figure 2A, 2D), suggesting that the absorbance maxima for standard curcuminoids and TPNP are about 420-430 nm, which agrees with published literature on curcumin’s absorbance maxima [44]. The absorbance-based method for quantifying total curcuminoids in aqueous media was found to have low sensitivity due to the colloidal nature of the samples, and should not be used for quality assurance or loading determinations of the API. Therefore, we adopted a fluorescence-based spectroscopic approach using 80% ethanol, which was subsequently validated by HPLC-DAD.

**Figure 2:**
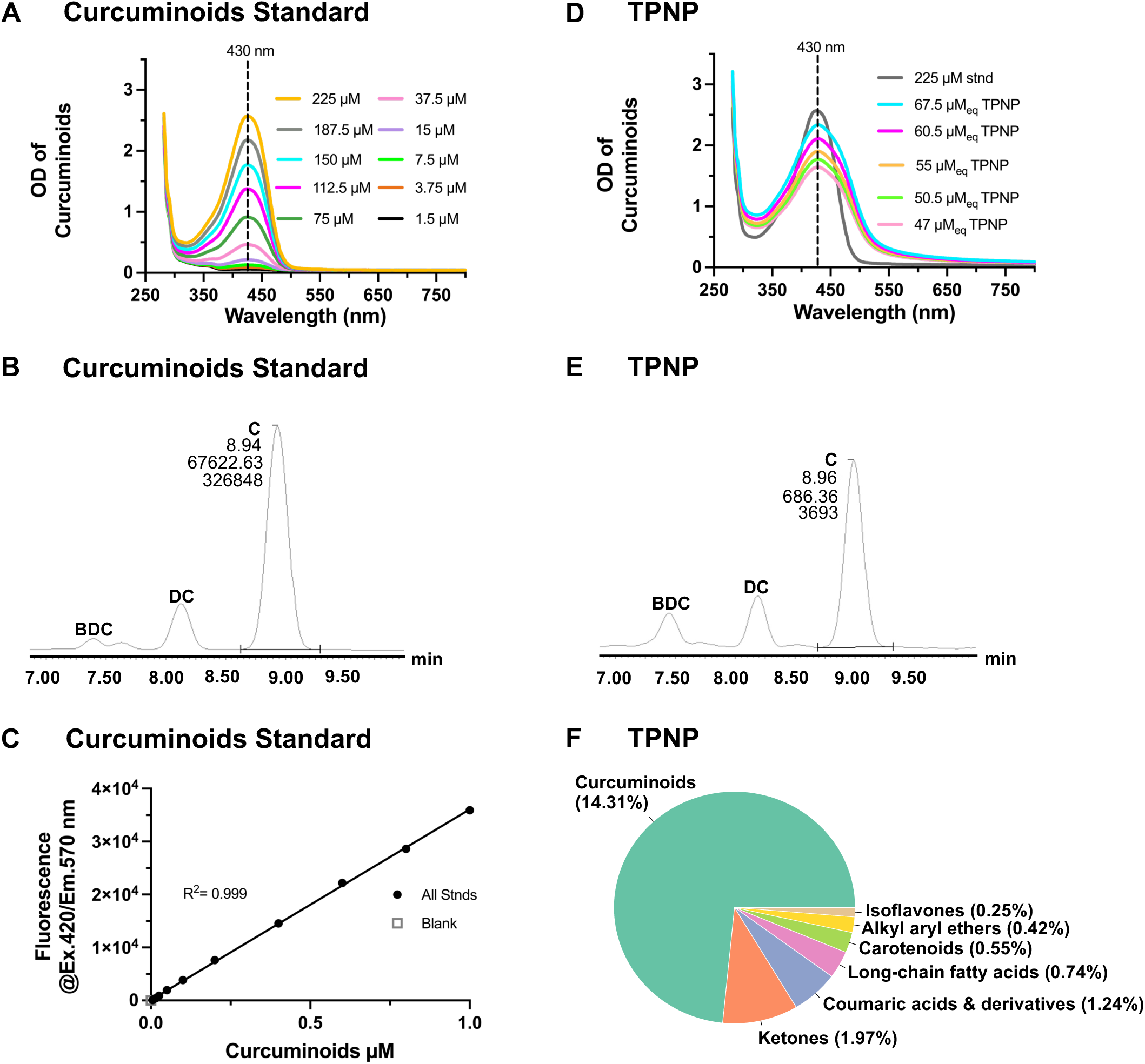
Characterization of curcuminoids reference standard (Sigma-Aldrich) showing (A) absorbance spectra of serial dilutions in water with a characteristic visible-range peak maximum near 430 nm, (B) mass-fraction distributions of individual curcuminoids—curcumin (C), desmethoxycurcumin (DC), and bisdemethoxycurcumin (BDC)—with values below the “C” peaks indicating retention time (min), peak area, and peak height (top to bottom), and (C) a fluorescence-based standard curve derived from serial dilutions enabling quantification of curcuminoid content via intrinsic fluorescence (Ex 420 nm/Em 570 nm). Comparative characterization of TPNP relative to curcuminoid reference standards showing (D) absorbance spectra of serial dilutions in water with a visible-range peak maximum near 430 nm, consistent with standard curcuminoids, (E) mass-fraction distributions of curcumin (C), desmethoxycurcumin (DC), and bisdemethoxycurcumin (BDC), and (F) Nano-LC–MS/MS-based chemo-ontological classification (ClassyFire Level-5) confirming that TPNP is predominantly composed of curcuminoids.

The rationale for using 80% ethanol as a diluent in fluorescent spectroscopy is to release all curcuminoids from TPNP and CSAP into a clear, non-colloidal solution for spectral measurements and to limit self-aggregation. For the fluorescence-based spectroscopy assay, the standard Sigma-Aldrich curcuminoid stock and TPNP were diluted in 80% ethanol followed by measurement with fluorescent excitation at 420 nm and emission capture at 570 nm. This selection was made because 420 nm serves as a common excitation wavelength for all three curcuminoids and it provides near-maximal absorbance for all three species, which is suitable for use in rapid fluorescence spectroscopy assays. Using 420 nm excitation and 570 nm emission, a fluorescence standard curve with high linearity (R^2^ = 0.999) was established (Figure 2C). This curve served as a validated, routine, rapid, and cost-effective assay for determining total curcuminoid concentration across TPNP batches. Validation relied upon the absorbance maxima of curcumin, desmethoxycurcumin, and bisdemethoxycurcumin, which were measured individually, yielding peaks at 429 nm, 423 nm, and 419 nm, respectively (Figures S2A, S2B and S2C). Standard HPLC-DAD mass-fraction distribution of the three individual curcuminoid mixed together (100 ppm each) is shown in Figure S2D; 1mM standard purified curcuminoids (Sigma-Aldrich) diluted in ethanol is composed of three curcuminoids (curcumin–71.1% by weight, desmethoxycurcumin–15.7% and bisdemethoxycurcumin–3.9%) and the absolute pharmaceutical equivalent curcuminoids’ concentration in TPNP was verified. Relative mass fraction distribution of three curcuminoids of the Sigma-Aldrich curcuminoids standard and TPNP are shown in Figure-2B and 2E, respectively. The pharmaceutical equivalent total curcuminoids loaded in TPNP was 2.03 mM.

### 3.5 Total metabolite analysis of TPNP

To identify the total molecular ontological fingerprint of TPNP, untargeted NanoLC–MS/MS profiling was performed to generate MS spectral data. Data structural annotation was performed using ClassyFire [41], an automated structure-based classification tool that integrates multiple biomolecular databases for natural products. The ClassyFire framework organized molecular entities into a hierarchical ontology: (i) Kingdom level: organic and inorganic compounds, (ii) SuperClass: 26 organic and 5 inorganic categories, (iii) Class: 764 structural categories, (iv) SubClass: 1,729 subdivisions encompassing >10,000 compounds, (v) Levels 5–11: finer resolution based on chemical weight, spanning 910 to 2 entities. Figure 2F shows distribution of the top seven categories of molecules by percentage of signal peak intensity areas after ClassyFire level 5 structural annotation, showing predominance of curcuminoids signals. Similarly, the top seven categories of molecules present in TPNP, based on ClassyFire SubClass structural annotation, are shown in Figure S3.

Using absorbance measurements when coupled with particle disruption using 80% ethanol, the pharmaceutical-equivalent curcuminoid concentration of TPNPs was determined to be 1.19 mM. In contrast, the fluorescence-based method yielded a concentration of 1.91 mM for experimental samples, compared with 2.03 mM as measured by HPLC–DAD. The close agreement between these values indicates that the fluorescence method, when coupled with 80% ethanol, accurately quantifies total curcuminoid content, with less than a 6% deviation from HPLC–DAD measurements. These findings support fluorescence (not absorbance) as a reliable approach for routine approximate quantification of total curcuminoids in TPNP samples. The dry weight of TPNP samples was determined to be 2877.6 µg / mL, containing 715 µg / mL total curcuminoids (sum of curcumin, desmethoxycurcumin and bisdemethoxycurcumin concentrations, as determined by HPLC-DAD), corresponding to a curcuminoid loading capacity of approximately 24.85% (w/w) in the TPNP samples. The particle concentration of TPNP was 9.58 x 10^10^ /mL. Based on these values, the average dry weight of a single TPNP was estimated to be approximately 30.29 fg (femtogram), of which ∼7.5 fg corresponds to curcuminoids; the remaining mass consists of compounds derived from the same turmeric rhizomes.

### 3.6 Direct antioxidant capacity of TPNP

Curcuminoids and other polyphenols are well known to exhibit potent antioxidant properties. Establishing a reference for this intrinsic ability is relevant to formulation quality assurance controls. To evaluate the antioxidant capacity of TPNPs, the FRAP assay–based on the reduction of ferric (Fe³⁺-TPTZ) to ferrous (Fe²⁺-TPTZ) ions–was performed on TPNPs and standard curcuminoids. TPTZ refers to the ligand scaffold that chelates iron, producing the characteristic, blue-colored complex in the FRAP assay. NAC (N-acetylcysteine) and Gallic Acid (GA) served as positive reference antioxidants in the assay. To calculate % antioxidant capacity, we used equimolar concentrations (62.5 µM) of antioxidants (curcuminoids standard, NAC and GA), where 62.5 µM of standard curcuminoids (Sigma-Aldrich) was used as a 100% antioxidant capacity reference (Figure 3A). The TPNP suspension medium or vehicle showed no antioxidant activity. The antioxidant capacity observed at 62.5 µM curcuminoid equivalent dose of TPNP is approximately 2.6 times greater than that of 62.5 µM standard curcuminoids (Figure-3A). This could be attributed to the presence of other bioactive antioxidant compounds derived from turmeric rhizome biomass, which accounts for approximately 75% of the total mass of TPNP. As previously shown in Figure 2F, NanoLC–MS/MS analysis of TPNP confirmed the presence of other metabolites, such as terpenoids, flavonoids, and carotenoids, which could contribute to higher antioxidant capacity.

**Figure 3:**
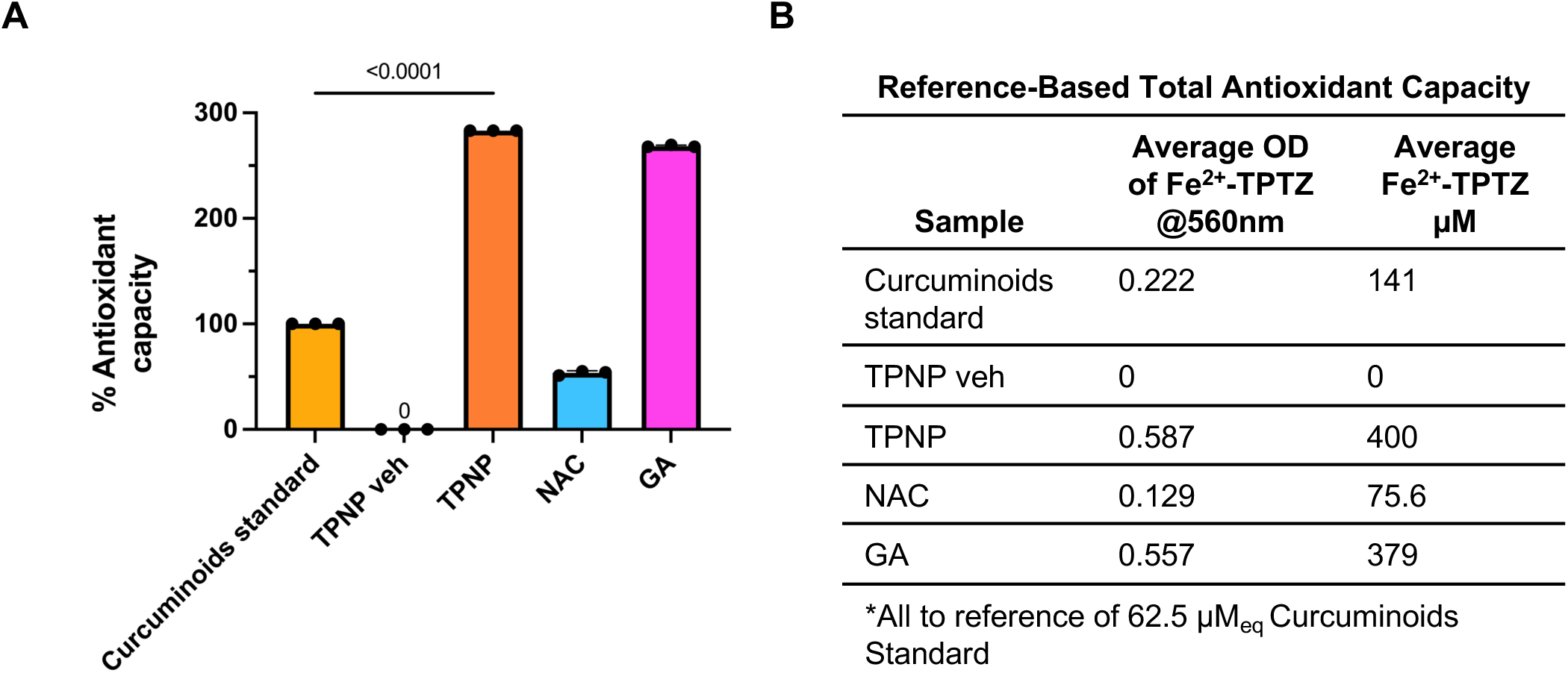
TPNP shows greater % antioxidant capacity compared to standard curcuminoids (Sigma-Aldrich). Ferric Reducing Antioxidant Power (FRAP), a colorimetric (blue) assay designed for the quantification and detection of Ferric Antioxidant Status was used to detect the (A) % antioxidant capacity at 62.5 µM equivalents of standard curcuminoids (Sigma), TPNP, N-Acetyl-L-Cysteine (NAC), and Gallic Acid (GA).

### 3.7 Dose tolerance of pharmaceutical equivalent concentrations of curcuminoids in CSAP and TPNP in THP-1 monocytes / macrophages

Tolerance testing is an important consideration to quality assurance controls and safety of product development of APIs. The LD_50_ of CSAP and TPNP was determined by exposing THP-1 monocytes and macrophages to equivalent concentrations of curcuminoids and performing a resazurin-based cell viability analysis (Figure-4). After 30 h exposure in monocytes, the LD_50_ of CSAP was approximately 9.24 µM, whereas that of TPNP was 30% higher, at 12.0 µM (Figure 4A, 4B). Similarly, THP-1 macrophages were exposed to varying doses of CSAP and TPNP, resulting in an LD_50_ of 10.3 µM for both treatments (Figure 4C, 4D).

**Figure 4:**
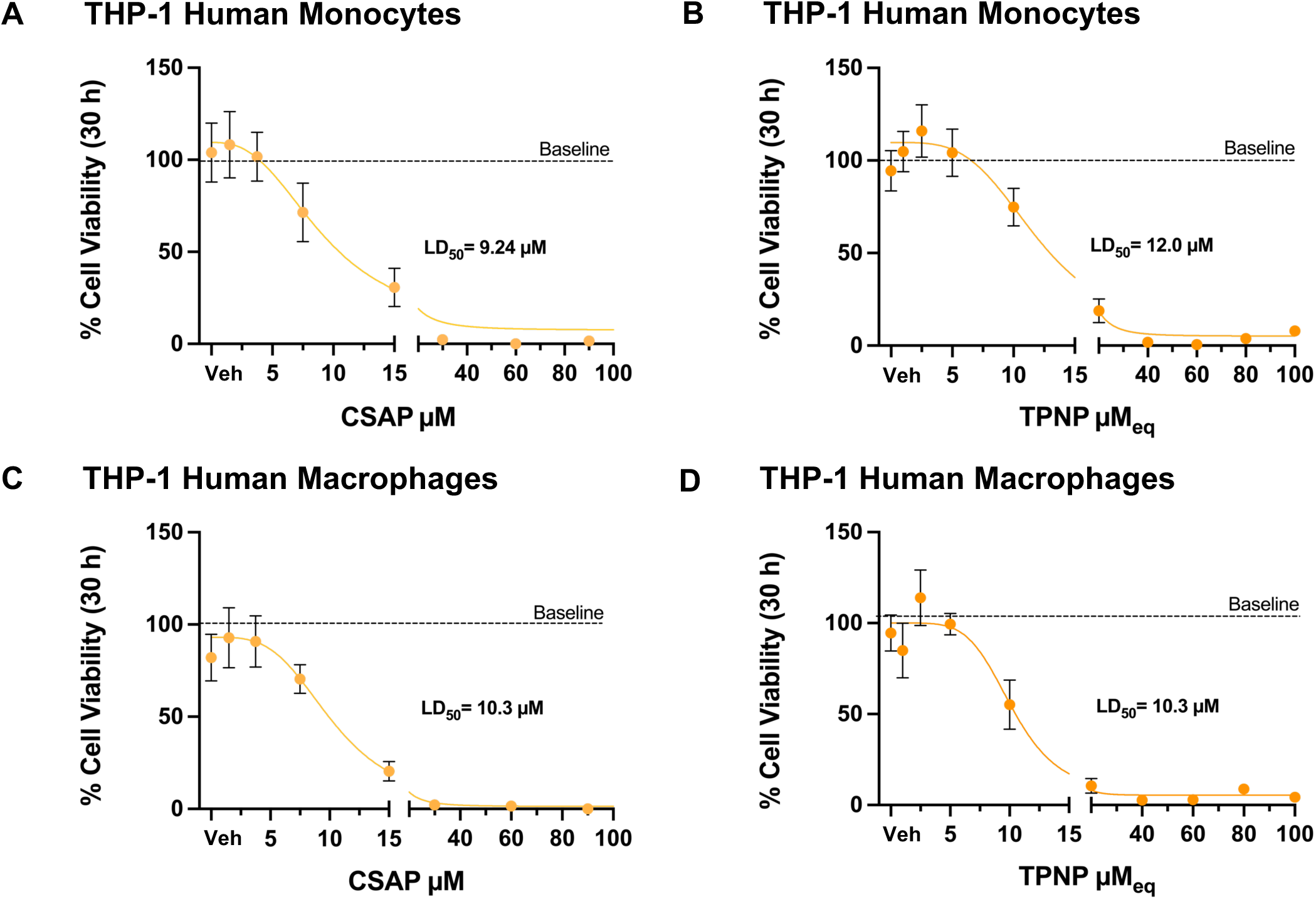
TPNP is slightly better tolerated than CSAP in monocytes, while both drugs show comparable tolerance in macrophages. Resazurin-based cell viability assay assessing dose-dependent tolerance of curcuminoids delivered as Curcuminoid Standard Spontaneous Aggregating Particles (CSAP) in monocytes (A) and macrophages (C), and as Turmeric-Phyto-NanoParticles (TPNP) in monocytes (B) and macrophages (D). Treatment concentrations were expressed as µM curcuminoids. Data was normalized to a media-only control. EtOH veh denotes the vehicle control for CSAP, whereas TPNP veh denotes the vehicle control for TPNP (PBS + EtOH).

#### 3.7.1. Dampening of LPS induced inflammation with treatments of TPNP in THP-1 macrophages

References to pertinent molecular mechanisms of action for minimal viable efficacy and comparability are essential for establishing quality assurance controls for nanoformulations used in medicinal contexts. To evaluate the cellular pharmacodynamic effects of TPNP and CSAP, we selected two cellular pharmacodynamic markers: pro-inflammatory cytokine TNF (measured by ELISA) and the anti-inflammatory enzyme HMOX1 (assessed by immunoblotting). THP-1 macrophages respond to LPS stimulation by rapidly releasing TNF in the extracellular media *in vitro*[42]. Curcuminoids are known to attenuate this response by downregulating TNF expression and secretion. The superior efficacy of a colloidal formulation (TPNP) compared to a reference standard (CSAP) was demonstrated by two strategies: either by achieving greater therapeutic effects at equivalent curcuminoid concentrations, or by matching or exceeding efficacy at a lower dose of the colloidal formulation with equivalent induction of the attributable mechanism of action, HMOX1. For the TNF investigation of efficacy, we applied the former approach to account for both the API’s intrinsic properties (antioxidant) and the indirect effects (mediated by HMOX1). THP-1 macrophages were pre-treated with a 5 µM equivalent dose of curcuminoids from TPNP or CSAP for 0.5 h, followed by 50 ng/mL LPS stimulation for a period of 1.5 h and 4 h. As shown in Figure 5A, TNF was released by these cells and detected in the conditioned media in 1.5 and 4 h post-LPS exposure. Treatment with 5 µM_eq_ TPNP or 5 µM CSAP on LPS-challenged macrophages reduced total secretion of TNF relative to media control (CSAP, p<0.0001; TPNP, p<0.0001–not shown on plot) at 1.5 h and 4 h. At 4 h, TPNP produced a markedly greater reduction in TNF secretion than CSAP (p<0.0001), with a similarly superior effect observed at 1.5 h (p = 0.0014); however, the early TPNP effect at 1.5 h is partially obscured by the TPNP vehicle response at this time point (p<0.0001; not shown on plot). No significant reduction in TNF release was observed when THP-1 monocytes were subjected to the same treatment and LPS regimens (data not shown).

**Figure 5:**
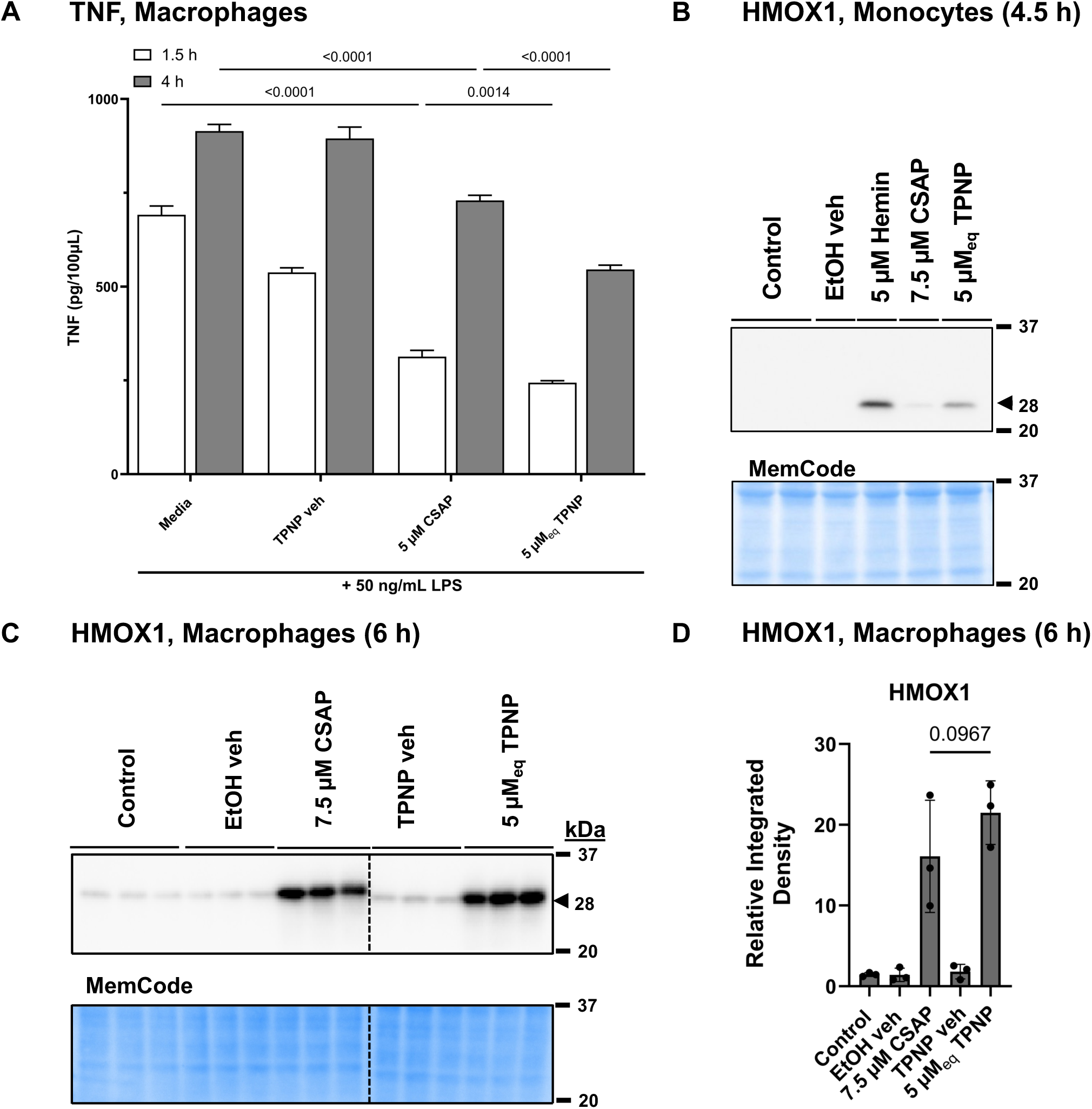
TPNP is superior at reducing TNF levels compared to CSAP at equivalent doses and is a more efficacious inducer of HMOX1. (A) TNF concentration measured in the conditioned media of macrophages treated with equal concentrations of TPNP and CSAP under the specified time and treatment conditions. (B) Representative immunoblot showing HMOX1 protein levels in THP-1 monocytes treated with TPNP (5.0 µM), CSAP (7.5 µM), vehicle control or media only. Hemin (5 µM) was included as a positive control for HMOX1 induction. (C,D) HMOX1 protein levels in THP-1 macrophages following treatment with the indicated drugs. Lower panels of (B) and (C) show total protein MemCode stain.

#### 3.7.2. HMOX1 protein levels in TPNP-treated THP-1 monocytes and macrophages

Increased protein levels of HMOX1 can promote anti-inflammatory and secondary antioxidant effects as heme catabolism by HMOX1 generates carbon monoxide and bilirubin, both of which possess cytoprotective and antioxidant properties that support wound healing in damaged tissues. Here, we sought to determine the equivalent HMOX1 induction with a lower curcuminoid concentration by TPNP than CSAP. HMOX1 protein levels were evaluated by immunoblotting CSAP or TPNP-treated THP-1 monocytes. Figure 5B shows a representative immunoblot of HMOX1 protein levels after 4.5 h exposure of 7.5 µM CSAP and 5 µM_eq_ TPNP in monocytes. To visualize the HMOX1 immunoblot signal, a higher amount of CSAP was required. No HMOX1 protein was observed in control treatments, but relatively higher protein levels were detected with 5 µM_eq_ TPNP compared to 7.5 µM CSAP. A 5 µM hemin treatment was included as a positive control for HMOX1 induction. Similarly, THP-1 macrophages were treated with 5 µM_eq_ TPNP and 7.5 µM CSAP for 6 h to achieve comparable induction of HMOX1 protein levels without adverse effect. TPNP induced equivalent HMOX1 in macrophages at a lower curcuminoid dose in comparison to a higher dose of CSAP (Figure 5C and 5D). Thus, TPNP exhibits a higher HMOX1 induction efficiency compared to CSAP. However, not all cell types are expected to tolerate high levels of HMOX1 induction. Yet, when exposed to these HMOX1 reference concentrations, AC16 human cardiomyocytes and hCMEC/D3 human blood–brain barrier (BBB) cells showed higher cell viability at 5 µM_eq_ TPNP than with 7.5 µM CSAP (Figure S4A, S4B), suggesting that TPNP was well tolerated. Based on HMOX1-equivalent doses determined from macrophage data (7.5 µM CSAP was comparable to 5 µM_eq_ TPNP), and HMOX1 referenced induction was well tolerated by TPNP in various cells including macrophages.

#### 3.7.3. Cellular bioavailability of CSAP and TPNP in THP-1 monocytes and macrophages

To assess whether the observed differences in HMOX1 protein levels and TNF suppression were attributable to differences in macrophage uptake kinetics between CSAP and TPNP, we conducted time- and dose-dependent FACS analysis. This approach leveraged the intrinsic autofluorescent nature of curcuminoids to quantify and compare their intracellular bioavailability. The internalization and accumulation of 5 µM_eq_ TPNP was first observed by confocal microscopy in THP-1 macrophages, where TPNP (detected via autofluorescence of curcumin captured at ex = 488 nm and em= 517 nm) localized in the nucleus and cytoplasm (Figure 6A). 5 µM_eq_ TPNP internalized rapidly in these cells, accumulating in 47.5% of cells (median fluorescent intensity= 527) within 0.5 h of exposure (Figure 6B(i), Table S1) and progressively accumulating in 95.3% of cells (median fluorescent intensity= 1271) at 16 h (Figure 6B(v), Table S1), whereas despite the higher particle availability of 7.5 µM CSAP it accumulated in only 8.05% of cells (median fluorescent intensity=277) within 0.5 h of exposure (Figure6B(i), Table S1) and reaching 28.8% of cells (median fluorescent intensity= 423) at 2 h of exposure (Figure6B(ii), Table S1), followed by progressively less curcuminoid positive cells from 4h-24h (Figure6B(iv-vi), Table S1).

**Figure 6:**
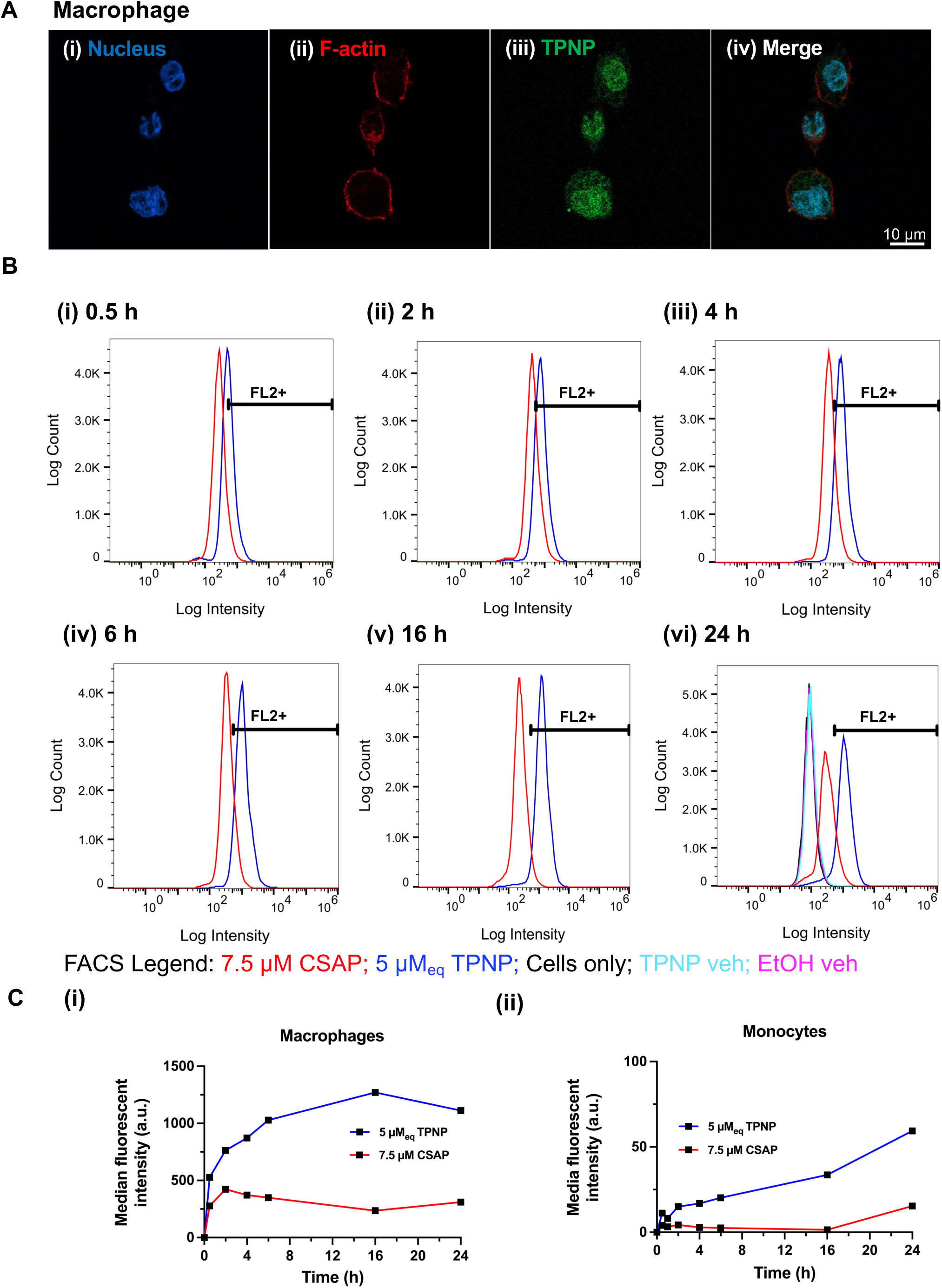
TPNP bioaccumulation is more rapid and cumulatively greater than CSAP in monocytes and macrophages. (A) Representative fluorescence microscopy images showing the subcellular distribution of TPNPs in THP-1 macrophages. (i) Nuclei stained with Hoechst (blue), (ii) F-actin stained with Phalloidin (red), (iii) TPNPs visualized via intrinsic green fluorescence, (iv) Merged images of three previous panels. (B) Representative time-course analysis of intracellular curcuminoid bioavailability in THP-1-derived macrophages treated with CSAP or TPNP. Fluorescence was assessed at 0.5 h, 2 h, 4 h 6 h, 16 h and 24 h time points. (C) Quantification of curcuminoid median fluorescent intensity over time in (i) macrophages and (ii) monocytes. Curcuminoid signal reflects intrinsic autofluorescence detected in the FL2+ channel.

Similarly, THP-1 monocytes were treated with pharmaceutical doses of 7.5 µM CSAP and 5 µM_eq_ TPNP for up to 24 h (Figure 6C(ii), Figure S5 and Table S2). FACS analysis demonstrated the accumulation/internalization of 5 µM_eq_ TPNP in monocytes reaching 16.9% cells (median fluorescent intensity= 739) after 4 h of exposure (Figure S5A(iii), Table S2) and highest at 59.4% of cells (median fluorescent intensity= 1397) after 24 h of exposure (Figure S5(vi), Table S2), whereas 7.5 µM CSAP dose accumulated to 2.85% (median fluorescent intensity= 313) in 4 h (FigureS5A(iii), Table S2) and highest about 15.4% of cells (median fluorescent intensity= 659) after 24h of exposure (Figure S5(vi), Table S2). These data indicate that tolerance and induction are not a byproduct of lower uptake, but rather show the opposite, with preferential phagocytosis and stable internalization of TPNP.

### 4.0 Discussion

TPNPs are bio-native, micelle-like nanoparticles (177 nm, −0.189mV) directly extracted from turmeric rhizomes and are distinct from plant-derived vesicles or exosomes. They contain 24.85% curcuminoids by mass, with the remainder comprising endogenous turmeric phytochemicals. In this study, additive-free TPNPs were compared with commercial extracted and solubilized curcumin, CSAP, which form nanoparticles spontaneously in cell culture media under surfactant- and carrier-free conditions and therefore provide a relevant physicochemical control. TPNPs and CSAPs exhibited submicron diameters by DLS, and TPNPs maintained colloidal homogeneity after lyophilization and rehydration. In the presence of fetal bovine serum (FBS), CSAPs decreased in size, consistent with protein-mediated stabilization, whereas TPNP size increased, potentially due to bio-corona formation. Curcuminoid content was confirmed by HPLC-DAD and fluorescence spectroscopy (coupled with EtOH for nanoparticle membrane disruption), and untargeted Nano-LC-MS/MS verified their complex phytochemical composition. In THP-1 monocytes and macrophages, both TPNP and CSAP induced dose-dependent toxicity, with slightly higher tolerance to TPNP in monocytes and comparable LD_50_ values in macrophages. Both TPNPs and CSAPs upregulated HMOX1, a cell stress marker and mediator of antioxidative and anti-inflammatory macrophage polarization, with more pronounced induction in macrophages than monocytes. Notably, TPNPs elicited greater HMOX1 upregulation at a lower dose. Flow cytometry using curcuminoid autofluorescence revealed significantly greater intracellular accumulation of TPNPs than CSAPs from 0.5 to 24 h, indicating more efficient cellular internalization despite their larger size and suggesting that compositional features, potential macrophage polarization state, or bio-corona interactions may be determining factors for bioavailability that particle size alone. Consistent with this enhanced uptake, TPNPs more effectively suppressed LPS-induced TNF secretion in THP-1 macrophages at equivalent curcuminoid doses, supporting superior anti-inflammatory efficacy and positioning TPNPs as a bioactive and safe alternative to synthetic nano-emulsions and lipid-based carriers for improving the cellular pharmacokinetics and pharmacodynamics of curcuminoids.

Others have evaluated the biophysical properties of natural delivery systems of curcumin. For example, curcumin-loaded sunflower seed protein isolate nanoparticles (SFPI-curcumin NPs) exhibited an average size of 194 ± 7 nm with a spherical and smooth surface[43]. These findings are consistent with this TPNP preparation, which exhibited a reported mean size of 177 ± 0.964 nm and a spherical morphology. By contrast, an alternative natural curcumin formulation—curcumin-loaded human serum albumin (HSA) nanoparticles—was reported to be approximately 200 nm in size, displaying a cube-like morphology with smooth surfaces, and a zeta potential of −10 mV[44]. In comparison, the TPNPs exhibited a near-neutral zeta potential, a difference that likely reflects fundamental distinctions in nanoparticle composition and surface chemistry. Curcumin-loaded soybean protein isolate–fucoidan nanocomposites exhibited irregular, non-uniform morphologies, in marked contrast to the highly uniform TPNP preparation observed here[45]. These differences likely originate from distinct formulation strategies: soybean protein isolate–fucoidan nanoparticles were produced via ultrasound processing and polymer self-assembly[45], whereas TPNPs are derived through extrusion of whole turmeric rhizomes to isolate bio-native, micelle-like structures, a process that better preserves spherical structural integrity rather than relying on the precise reassembly of discrete soybean and fucoidan building blocks.

There are limited studies that explicitly acknowledge and characterize the spontaneous formation of curcumin self-assembled particles (CSAPs) in cell culture media. Notably, Kang *et al.* reported that the average particle size of curcumin increased progressively across mediums, measuring approximately 173, 590, and 903 nm in distilled water, 1x PBS, and RPMI-1640 medium alone, respectively[40]. In contrast to those findings, our results show that CSAP exhibited a mean particle size of 410.1 ± 8.7 nm in RPMI-1640 alone, whereas a substantially larger size of 1088.0 ± 54.0 nm was observed in 0.1x PBS; this contrasts with the 590 nm reported in 1x PBS by Kang *et al*. This discrepancy in PBS-based measurements may be attributed to differences in buffer ionic strength and buffering capacity between 1x PBS and the diluted 0.1x PBS used in this study. Reducing ionic strength can diminish electrostatic screening and destabilize curcuminoid aggregates, thereby promoting particle–particle interactions and secondary aggregation, yielding larger apparent particle sizes. Collectively, these findings highlight the sensitivity of curcumin formations (CSAP) to subtle changes in solution chemistry and underscore the importance of buffer composition when interpreting nanoparticle sizing data.

Accurate quantification of curcuminoids is highly method-dependent, with detector sensitivity and spectral optimization being paramount in quantifying loading of curcuminoids into nanoparticles. One study defined the fluorescence excitation and emission maxima of curcumin at 485 and 528 nm, respectively, yielding a highly linear curcuminoid standard curve (R² = 0.998)[46]. Although our excitation (420 nm) and emission (570 nm) wavelengths differ modestly, this difference likely reflects variations in solvent composition (EtOH vs DMSO), spectral shifts, and possible unaccounted-for background effects. In that prior study, solid lipid nanoparticles (SLNs; 542.40 nm, 53.50 mV) and chitosan nanoparticles (Chi-NPs; 537.10 nm, −17.70 mV) were quantified using both spectrofluorimetric and HPLC methods. For Chi-NPs, fluorescence quantified 2.12 ± 0.182 mg/mL compared to 3.36 ± 0.227 mg/mL curcumin by HPLC, representing a ∼37% lower value by fluorescence. Similarly, for SLNs, fluorescence measured 2.48 ± 0.12 mg/mL versus 3.17 ± 0.141 mg/mL curcumin by HPLC, corresponding to a ∼22% lower concentration[46]. Our findings demonstrate that curcuminoid quantification in TPNPs deviated by less than 6% between fluorescence and HPLC analyses, indicating strong agreement between the two methods. This close concordance may be attributed, in part, to the optimized selection of excitation and emission wavelengths in the spectrofluorimetric assay, together with the use of 80% EtOH to disrupt nanoparticle membranes and fully solubilize curcuminoids, thereby improving analyte recovery, enhancing fluorescence signal accuracy, and reducing matrix-dependent variability. These differences underscore the higher sensitivity and recovery afforded by HPLC methods while highlighting the utility of fluorescence coupled with EtOH as a rapid and inexpensive quantification approach for nanoparticle-encapsulated curcuminoids.

Liakopoulou *et al.* demonstrated that curcumin-loaded solid lipid nanoparticles (SLN.CUR), nanostructured lipid carriers (NLC.CUR), and nanoemulsions (NE.CUR) exhibited significantly greater total antioxidant capacity than soluble curcumin, as assessed by the FRAP assay[47]. Consistent results have also been reported for curcumin-loaded sunflower seed protein isolate nanoparticles (SFPI–curcumin NPs), which similarly outperformed soluble curcumin in FRAP-based antioxidant measurements[43]. Collectively, these findings align with the present observations, wherein TPNPs exhibited superior antioxidant performance compared to soluble curcuminoids. This also affirms the intrinsic benefits purported for nanoformulated curcumin, which are not limited to variations in absorption or cellular uptake (particularly relevant to orally formulated products).

Understanding curcumin’s cellular tolerance in both free and nanoparticle formulations is critical for optimizing therapeutic efficacy while minimizing adverse effects across diverse cell types. Direct head-to-head dose–response comparisons between nanocurcumin and free curcumin at equivalent curcuminoid concentrations could be insufficient and result in unfair comparisons compared to the indirect pleiotropic effects mediated by mechanisms of action like HMOX1. Evidence particularly in macrophages, a main target of anti-inflammation, remain limited, one macrophage study evaluated the anti-inflammatory, antioxidative, and anti-angiogenic effects of nanocurcumin in RAW 264.7 mouse macrophages, identified 5 µM_eq_ as a safe dose at 24 h post dose–response analysis[48]. Similarly, we present a dose–response assessment here to determine therapeutic range and identify 5 µM_eq_ TPNPs as relatively safe *in vitro across various human cell types*. Another independent study showed that in MCF7 human epithelial-like breast cancer cells, the IC_50_ of curcumin-loaded micelles was 13.9 µg/mL or 38.0 µM, whereas free curcumin was non-toxic across all tested concentrations (2.3–600 µg/mL or 6.2 µM-1.6 mM) after 72 h exposure[49]. This suggests uncertainty unless uptake is accounted for and formulation chemistry is shelf stable during testing. In other cell types, higher tested nanocurcumin or curcumin doses have been employed—up to 5 µg/mL or 14 µM_eq_ nanocurcumin in Human Umbilical Vein Endothelial Cells (HUVECs)[50] or 10 µM curcumin in RLE-6TN rat epithelial cells for 24 h[51]—yet the latter study lacked a dose–response assessment. Notably, these concentrations exceed our maximum tested dose of 5 µM_eq_ TPNPs or 7.5 µM CSAP, possibly due to our slightly longer 30 h incubation period. In other cell types, curcumin-loaded tannic acid–poloxamer (Cur-TA-poloxamer) nanoparticles have been reported to exhibit lower cellular tolerance than soluble curcumin in MDA-MB-231 human mesenchymal-like breast cancer cells (IC_50_; 10.6 vs 18.7 µM), PANC-1 human pancreatic cells (IC_50_; 9.2 vs 28.2 µM), and U-87 human glioblastoma cells (IC_50_; 14.7 vs 28.2 µM) after 72 h exposure[52]. The IC_50_ of Cur-TA-poloxamer in MDA-MB-231 mesenchymal-like human breast cancer cells is consistent with the LD_50_ of 10.3 µM observed in macrophages exposed to TPNPs in our study, despite the shorter duration of drug exposure. Collectively, these findings highlight the importance of dose–response characterization for optimizing efficacy and safety across both phagocytic and non-phagocytic cell types, and suggest that nanoparticle formulations of curcumin can exhibit greater cytotoxicity than soluble curcumin, likely reflecting enhanced intracellular delivery and accumulation afforded by the nanoformulation. Moreover, the selection of cancerous vs non-cancerous, or immortalized vs primary, cells is a potential confounder *in vitro* studies when translating efficacy in vivo or clinically. The use of cell organoids and non-cancerous or primary human cells may be advisable before formally advancing to investigational new drug or preclinical outcome-based evaluations.

Nanoformulations fundamentally enhance the cellular pharmacokinetics and pharmacodynamics of curcumin, resulting in superior intracellular accumulation and downstream anti-inflammatory and anti-oxidant effects compared with free curcumin across both phagocytic and non-phagocytic cell types. Consistent with our observations, a previous study reported significantly higher mean fluorescence intensity for 10 µM_eq_ curcumin encapsulated in tannic acid–poloxamer (Cur-TA-poloxamer; 193 nm) compared with 10 µM free curcumin following 4 h of incubation in both epithelial-like MCF-7 cells (mean fluorescence intensity: 5990 vs 1000) and mesenchymal-like MDA-MB-231 breast cancer cells (mean fluorescence intensity: 5500 vs 3350) by flow cytometry, indicating enhanced cellular pharmacokinetics not only in macrophages, as demonstrated here, but also in non-phagocytic cell types[52]. Confocal microscopy from an independent study further corroborated enhanced intracellular uptake of the *in situ* polymeric PLGA curcumin-loaded nanoparticles (ISCurNP; 208.25 nm), demonstrating that curcumin nanoparticles promote greater intracellular accumulation compared to free curcumin in RAW 264.7 mouse derived macrophages after 3 h exposure[53]. Another study reported that curcumin-loaded polydopamine–fructose (PCF) nanoparticles, with PCF3 being the smallest (106 nm) and PCF144 being the largest (134 nm), exhibited decreased cellular uptake with increasing particle size after 6 h of exposure to RAW 264.7 mouse macrophages, MCF-7 human epithelial-like breast cancer cells and bovine aortic endothelial cells (BAECs) by flow cytometry[54]. Importantly, neither of these studies accounted for the autoformation of curcumin aggregates in cell culture media. Yet, the hydrodynamic size of Cur-TA-poloxamer nanoparticles (193 nm) and ISCurNP (208.25 nm) are comparable to TPNP (223.8 nm) under cell culture conditions, where Cur-TA-poloxamer and ISCurNP exhibited more rapid and greater cellular bioaccumulation than free curcumin. Although PCF144 was larger than PCF3, it did not lead to increased intracellular uptake, further demonstrating that increasing nanoparticle size does not inherently improve cellular internalization. Collectively, these findings indicate that the markedly enhanced macrophage uptake of TPNP observed in our study is driven primarily by its compositional and surface characteristics rather than nanoparticle size alone. Sarawi *et al*. showed that lung tissues collected from CuSO_4_-intoxicated rats (i.e., CuSO_4_ triggers lung injury in rats by causing oxidative stress) treated for 7 days with 80[mg/kg]_eq_ nanocurcumin (liposomal encapsulated curcumin; nCurc) exhibited higher induction of HMOX1 compared to 80[mg/kg] curcumin[55]. RAW 264.7 mouse macrophages exposed to Curcumin-loaded lipid-poly(lactic-co-glycolic acid hybrid microparticles (Cur@MPs) or free Cur for 24h followed by to LPS (1000ng/mL or 1µg/mL) for 24 h exhibited greater attenuation of TNF with Cur@MPs compared to free curcumin[48]. Another study reported that exposure to polyvinylpyrrolidone-coated curcumin nanoparticles (PVP coated Cur-NPs; 10 µM_eq_) for 12–72 h significantly attenuated TNF secretion in NR8383 rat alveolar macrophages following LPS (5 µg/mL) stimulation[56]. Taken together, curcumin nanoparticles are likely internalized more efficiently than curcumin reference standards both *in vitro* and *in vivo*, enabling greater intracellular delivery of curcumin and thereby promoting stronger induction of the pleiotropic mechanisms of action, like HMOX1, to achieve greater suppression of pro-inflammatory TNF following LPS stimulation. These effects are likely driven by the nanoparticles’ distinct compositional and surface characteristics, as well as the probability for opsonization, corona formation, and other interactions with biological media that require further study.

A key limitation of this study is the absence of a head-to-head comparison of 5/ 7.5 µM_eq_ TPNPs and of 5/7.5 µM CSAPs in the FRAP assay. This may help elucidate how equivalent doses of TPNP and CSAP perform in a direct chemical antioxidant assay and complement the *in vitro* assays assessing their biological effects. However, matching the curcuminoid concentration between TPNPs and CSAPs was straightforward—and CSAPs served as an appropriate control given their nanometric size, additive-free formulation, and comparable curcuminoid composition. Precisely matching particle size between the two preparations was not feasible, and the goal was to evaluate direct antioxidant effects. Achieving strict size equivalence by nano sieving might also enable clearer delineation of the extent to which the larger size of TPNPs, relative to CSAPs, contributes to enhanced macrophage uptake and, in turn, impacts therapeutic efficacy. These methods have not yet been developed or optimized but could in the future be used to determine relevance.

HPLC-DAD and ethanol disruption coupled with fluorescence are robust methods for quantifying API in delivery systems such as TPNPs, whereas absorbance measurements are comparatively unreliable. This may explain some of the prolonged discrepancies in results in the field when evaluating curcumin and its derivatives. Importantly, these results highlight that evaluating the colloidal API alone in simplified chemical assays and *in vitro* systems is insufficient to predict therapeutic potential, and that both the API and mechanism of action—here reflected by HMOX1 induction and TNF suppression—must be considered in parallel, or risk making erroneous attribution errors to both safety and efficacy. Using an direct antioxidant assay, equimolar comparisons between TPNPs and a reference curcuminoid standard isolated the contribution of the API itself, while head-to-head testing of TPNPs versus CSAPs at matched curcuminoid concentrations assessed the impact of colloidal architecture on cell tolerance and effective/biologically-equivalent dose. Flow cytometry further served as an API-sensitive assay, in which TPNPs consistently outperformed CSAPs in cellular uptake. Together, HMOX1 and TNF functioned as surrogate biological readouts demonstrating that TPNPs would likely achieve greater therapeutic effects than CSAPs at the same curcuminoid dose or maintain equivalent efficacy at lower colloidal doses, underscoring their therapeutic potential. The added benefit of shelf stability and ability to be lyophilized and rehydrated make utility in commercial development superior to current practices and methods of preparation of curcumin formulation for natural product or medicinal use. Although several studies have compared the attenuation of TNF signaling by nano-curcumin versus free curcumin *in vitro*, to our knowledge no published study has directly performed a head-to-head comparison of HMOX1 protein induction following treatment with nano-encapsulated versus soluble curcuminoids *in vitro*. Moreover, to our knowledge, this study is one of the few studies to explicitly consider the collective contribution of all curcuminoids when encapsulated within a natural nanoparticle formulation, thereby providing a more comprehensive and mechanistically accurate evaluation of curcuminoid-mediated antioxidative and anti-inflammatory effects through HMOX1 induction and TNF suppression. This distinction is critical, as the individual curcuminoid derivatives—desmethoxycurcumin and bisdemethoxycurcumin—have each been shown to exert independent anti-inflammatory and anti-oxidative effects[57][58][59].

The translatability of curcumin natural products and medicinal products to human benefit has been limited by intrinsic and unaccounted-for formulation effects on stability and by chemical and biological analytical methods. Here, we have established that an all-natural extraction and formulation process that generates curcumin nanoparticles directly from turmeric root, without reliance on synthetic excipient-based building blocks, yields nanoparticles that exhibit comparable, if not superior, pharmacokinetic/dynamic effects in human cells. Moreover, we advance the quality and control evaluation processes that should be minimally undertaken to evaluate such formulations against reference standards, accounting for chemical, biophysical, and biological effects that may involve direct and indirect mechanisms of action, thereby enabling fairer comparisons and advancing nanoproduct research and development.

## Acknowledgement

This research is supported by Natural Science and Engineering Research Council (NSERC), Research New Brunswick(RNB), Dalhousie University - Chesley Family Research Fund (to KRB), New Brunswick Innovation Foundation, National Research Council of Canada, and Mitacs Canada (to KRB and AG). The authors acknowledge the analytical and instrument support from the New Brunswick Regional Productivity Council.

## Disclosures

AG is a co-founder and holder of equity positions in Pividl Biosciences Inc., with process and composition of matter intellectual property related to the TPNP nanoformulation evaluated in this report. KRB holds a majority equity position in NB-BioMatrix Inc. and is a medical scientific advisor to Pividl Biosciences Inc. with equity options. Pividl Biosciences Inc. provided arm’s length trainee funds through Mitacs Canada Scholarships received in part by BCH and KRDW. No other authors have relevant disclosures or conflicts of interest to report at the time of this report.

---

**Figure S1: FBS in cell culture media increases the average hydrodynamic size of TPNPs, while it stabilizes and reduces the average hydrodynamic size of CSAP.** Dynamic Light Scattering (DLS) was used to assess the particle size distribution of CSAP and TPNP in various aqueous environments. (A-D) Size distribution profiles of CSAP in RPMI-1640 media with 0.5% FBS; PDI= 0.146 (A), RPMI-1640 media without FBS; PDI= 0.260 (B), 0.1x PBS with 0.5% FBS; PDI= 0.198 (C), CSAP in 0.1x PBS without FBS; PDI= 0.244 (D). (E-F) Size distribution profiles of TPNP in RPMI-1640 media with 0.5% FBS; PDI=0.121 (E) and without FBS; PDI=0.04 (F). DLS measurements reflect hydrodynamic diameter (d.nm) by intensity. All x-axis values are represented on log10 scale.

**Figure S2: Curcuminoid species exhibit distinct HPLC-DAD retention times and comparable absorbance maxima.** Representative HPLC-DAD absorbance spectra of 100 ppm standards for individual curcuminoids: (A) curcumin, (B) desmethoxycurcumin and (C) bisdemethoxycurcumin. All three species show strong absorbance in the visible range, with peak maxima ranging between 420nm-430nm. (D) Representative HPLC chromatogram of a curcuminoid mixture (100 ppm total) showing mass fraction distributions of curcumin (C), desmethoxycurcumin (DC) and bisdemethoxycurcumin (BDC). The numbers on the “C” peaks from top to bottom correspond to retention time (in min), peak area and peak height, respectively).

**Figure S3:** Annotated molecule categories of TPNP using nano-LC-MS/MS data for chemo-ontology ClassyFire SubClass, where Sesquiterpenoids are the major subclass present in the formulation.

**Figure S4: CSAP but not TPNP has lower tolerance at HMOX1 equivalent efficacy dose in AC16 human cardiomyocytes and hCMEC/D3 human BBB cells.** Treatment with 7.5 µM CSAP and 5 µM_eq_ TPNP in (A) AC16 human cardiomyocytes and (B) hCMEC/D3 human BBB cells.

**Figure S5: TPNP demonstrates enhanced cellular uptake compared to CSAP in THP-1 monocytes.** Representative FACS plots showing comparative intracellular bioavailability kinetics of CSAP and TPNP in THP-1 monocytes at 0.5 h, 2 h, 4 h 6 h, 16 h and 24 h time points. All signals detected in the FL2+ channel (curcuminoid positive).

**Table S1:** Table summarizing macrophage FACS data of percent of gated cells in the FL2+ channel (curcuminoid positive) with indicated time of treatments of CSAP and TPNP.

**Table S2:** Table summarizing monocyte FACS data of percent of gated cells in the FL2+ channel (curcuminoid positive) with indicated time of treatments of CSAP and TPNP.

## Notes

### Competing Interest Statement

The authors have declared no competing interest.

### Summary of Updates

To reorganize some text, expand the discussion.

## References

[1] W.-W. Tian et al., “Curcuma Longa (turmeric): from traditional applications to modern plant medicine research hotspots,” Chin. Med., vol. 20, no. 1, p. 76, May 2025, doi: 10.1186/s13020-025-01115-z.

[2] S. W. Ryter, “Heme Oxgenase-1, a Cardinal Modulator of Regulated Cell Death and Inflammation,” Cells, vol. 10, no. 3, p. 515, Feb. 2021, doi: 10.3390/cells10030515.

[3] V. Vijayan, F. A. D. T. G. Wagener, and S. Immenschuh, “The macrophage heme-heme oxygenase-1 system and its role in inflammation,” Biochem. Pharmacol., vol. 153, pp. 159–167, Jul. 2018, doi: 10.1016/j.bcp.2018.02.010.

[4] M. Zhang et al., “Myeloid HO-1 modulates macrophage polarization and protects against ischemia-reperfusion injury,” JCI Insight, vol. 3, no. 19, p. e120596, doi: 10.1172/jci.insight.120596.

[5] A. Sahebkar, A. F. G. Cicero, L. E. Simental-Mendía, B. B. Aggarwal, and S. C. Gupta, “Curcumin downregulates human tumor necrosis factor-α levels: A systematic review and meta-analysis ofrandomized controlled trials,” Pharmacol. Res., vol. 107, pp. 234–242, May 2016, doi: 10.1016/j.phrs.2016.03.026.

[6] Y. Niu et al., “Dietary Curcumin Supplementation Increases Antioxidant Capacity, Upregulates Nrf2 and Hmox1 Levels in the Liver of Piglet Model with Intrauterine Growth Retardation,” Nutrients, vol. 11, no. 12, p. 2978, Dec. 2019, doi: 10.3390/nu11122978.

[7] Z. Lin et al., “Bioinformatics and validation reveal the potential target of curcumin in the treatment of diabetic peripheral neuropathy,” Neuropharmacology, vol. 260, p. 110131, Dec. 2024, doi: 10.1016/j.neuropharm.2024.110131.

[8] T. Kushida, G. Li Volti, S. Quan, A. Goodman, and N. G. Abraham, “Role of human heme oxygenase-1 in attenuating TNF-alpha-mediated inflammation injury in endothelial cells,” J. Cell. Biochem., vol. 87, no. 4, pp. 377–385, 2002, doi: 10.1002/jcb.10316.

[9] U. Saqib, S. Sarkar, K. Suk, O. Mohammad, M. S. Baig, and R. Savai, “Phytochemicals as modulators of M1-M2 macrophages in inflammation,” Oncotarget, vol. 9, no. 25, pp. 17937–17950, Apr. 2018, doi: 10.18632/oncotarget.24788.

[10] W. Jin, B. O. A. Botchway, and X. Liu, “Curcumin Can Activate the Nrf2/HO-1 Signaling Pathway and Scavenge Free Radicals in Spinal Cord Injury Treatment,” Neurorehabil. Neural Repair, vol. 35, no. 7, pp. 576–584, Jul. 2021, doi: 10.1177/15459683211011232.

[11] Y. Zhong, J. Feng, Z. Fan, and J. Li, “Curcumin increases cholesterol efflux via heme oxygenase-1-mediated ABCA1 and SR-BI expression in macrophages,” Mol. Med. Rep., vol. 17, no. 4, pp. 6138–6143, Apr. 2018, doi: 10.3892/mmr.2018.8577.

[12] S. Shao, X. Ye, W. Su, and Y. Wang, “Curcumin alleviates Alzheimer’s disease by inhibiting inflammatory response, oxidative stress and activating the AMPK pathway,” J. Chem. Neuroanat., vol. 134, p. 102363, Dec. 2023, doi: 10.1016/j.jchemneu.2023.102363.

[13] M. R. Islam et al., “Targeted therapies of curcumin focus on its therapeutic benefits in cancers and human health: Molecular signaling pathway-based approaches and future perspectives,” Biomed. Pharmacother. Biomedecine Pharmacother., vol. 170, p. 116034, Jan. 2024, doi: 10.1016/j.biopha.2023.116034.

[14] E. Rapti, T. Adamantidi, P. Efthymiopoulos, G. Z. Kyzas, and A. Tsoupras, “Potential Applications of the Anti-Inflammatory, Antithrombotic and Antioxidant Health-Promoting Properties of Curcumin: A Critical Review,” Nutraceuticals, vol. 4, no. 4, pp. 562–595, Oct. 2024, doi: 10.3390/nutraceuticals4040031.

[15] K. M. Nelson, J. L. Dahlin, J. Bisson, J. Graham, G. F. Pauli, and M. A. Walters, “The Essential Medicinal Chemistry of Curcumin,” J. Med. Chem., vol. 60, no. 5, pp. 1620–1637, Mar. 2017, doi: 10.1021/acs.jmedchem.6b00975.

[16] S. J. Stohs, O. Chen, S. D. Ray, J. Ji, L. R. Bucci, and H. G. Preuss, “Highly Bioavailable Forms of Curcumin and Promising Avenues for Curcumin-Based Research and Application: A Review,” Molecules, vol. 25, no. 6, p. 1397, Mar. 2020, doi: 10.3390/molecules25061397.

[17] Y. Zhu, J. Ye, and Q. Zhang, “Self-emulsifying Drug Delivery System Improve Oral Bioavailability: Role of Excipients and Physico-chemical Characterization,” Pharm. Nanotechnol., vol. 8, no. 4, pp. 290–301, 2020, doi: 10.2174/2211738508666200811104240.

[18] H. H. Tayeb and F. Sainsbury, “Nanoemulsions in drug delivery: formulation to medical application,” Nanomed., vol. 13, no. 19, pp. 2507–2525, Oct. 2018, doi: 10.2217/nnm-2018-0088.

[19] S. Meirinho, M. Rodrigues, A. O. Santos, A. Falcão, and G. Alves, “Self-Emulsifying Drug Delivery Systems: An Alternative Approach to Improve Brain Bioavailability of Poorly Water-Soluble Drugs through Intranasal Administration,” Pharmaceutics, vol. 14, no. 7, p. 1487, Jul. 2022, doi: 10.3390/pharmaceutics14071487.

[20] L. Miclotte et al., “Dietary Emulsifiers Alter Composition and Activity of the Human Gut Microbiota in vitro, Irrespective of Chemical or Natural Emulsifier Origin,” Front. Microbiol., vol. 11, p. 577474, 2020, doi: 10.3389/fmicb.2020.577474.

[21] S. Naimi, E. Viennois, A. T. Gewirtz, and B. Chassaing, “Direct impact of commonly used dietary emulsifiers on human gut microbiota,” Microbiome, vol. 9, p. 66, Mar. 2021, doi: 10.1186/s40168-020-00996-6.

[22] S. Eswar, B. Rajagopalan, K. Ete, and S. Nageswara Rao Gattem, “Serum Tumor Necrosis Factor Alpha (TNF-α) Levels in Obese and Overweight Adults: Correlations With Metabolic Syndrome and Inflammatory Markers,” Cureus, vol. 16, no. 7, p. e64619, Jul. 2024, doi: 10.7759/cureus.64619.

[23] M. De Siena et al., “Food Emulsifiers and Metabolic Syndrome: The Role of the Gut Microbiota,” Foods, vol. 11, no. 15, p. 2205, Jul. 2022, doi: 10.3390/foods11152205.

[24] A. S. Bancil, A. M. Sandall, M. Rossi, B. Chassaing, J. O. Lindsay, and K. Whelan, “Food Additive Emulsifiers and Their Impact on Gut Microbiome, Permeability, and Inflammation: Mechanistic Insights in Inflammatory Bowel Disease,” J. Crohns Colitis, vol. 15, no. 6, pp. 1068–1079, Jun. 2021, doi: 10.1093/ecco-jcc/jjaa254.

[25] Y.-B. Tan et al., “Effect of Ionic and Non-Ionic Surfactants on the Pasting Characteristics and Digestive Properties of Regular and Frozen Starch for Oral Delivery,” Foods, vol. 11, no. 21, Oct. 2022, doi: 10.3390/foods11213395.

[26] B. Chassaing et al., “Dietary emulsifiers impact the mouse gut microbiota promoting colitis and metabolic syndrome,” Nature, vol. 519, no. 7541, pp. 92–96, Mar. 2015, doi: 10.1038/nature14232.

[27] M. Eskandani, H. Hamishehkar, and J. Ezzati Nazhad Dolatabadi, “Cyto/Genotoxicity study of polyoxyethylene (20) sorbitan monolaurate (tween 20),” DNA Cell Biol., vol. 32, no. 9, pp. 498–503, Sep. 2013, doi: 10.1089/dna.2013.2059.

[28] C. Kriegel, M. Festag, R. S. K. Kishore, D. Roethlisberger, and G. Schmitt, “Pediatric Safety of Polysorbates in Drug Formulations,” Children, vol. 7, no. 1, p. 1, Dec. 2019, doi: 10.3390/children7010001.

[29] F. Lindenberg, F. Sichel, M. Lechevrel, R. Respaud, and G. Saint-Lorant, “Evaluation of Lung Cell Toxicity of Surfactants for Inhalation Route,” J. Toxicol. Risk Assess., vol. 5, May 2019, doi: 10.23937/2572-4061.1510022.

[30] B. Arechabala, C. Coiffard, P. Rivalland, L. J. Coiffard, and Y. de Roeck-Holtzhauer, “Comparison of cytotoxicity of various surfactants tested on normal human fibroblast cultures using the neutral red test, MTT assay and LDH release,” J. Appl. Toxicol. JAT, vol. 19, no. 3, pp. 163–165, 1999, doi: 10.1002/(sici)1099-1263(199905/06)19:3%3C163::aid-jat561%3E3.0.co;2-h.

[31] A. M. Silva et al., “Soft Cationic Nanoparticles for Drug Delivery: Production and Cytotoxicity of Solid Lipid Nanoparticles (SLNs),” Appl. Sci., vol. 9, no. 20, Oct. 2019, doi: 10.3390/app9204438.

[32] N. Vlachy, D. Touraud, J. Heilmann, and W. Kunz, “Determining the cytotoxicity of catanionic surfactant mixtures on HeLa cells,” Colloids Surf. B Biointerfaces, vol. 70, no. 2, pp. 278–280, May 2009, doi: 10.1016/j.colsurfb.2008.12.038.

[33] H. T. Lam, B. Le-Vinh, T. N. Q. Phan, and A. Bernkop-Schnürch, “Self-emulsifying drug delivery systems and cationic surfactants: do they potentiate each other in cytotoxicity,” J. Pharm. Pharmacol., vol. 71, no. 2, pp. 156–166, Feb. 2019, doi: 10.1111/jphp.13021.

[34] L. Gong et al., “Effect of polyethylene glycol on polysaccharides: From molecular modification, composite matrixes, synergetic properties to embeddable application in food fields,” Carbohydr. Polym., vol. 327, p. 121647, Mar. 2024, doi: 10.1016/j.carbpol.2023.121647.

[35] L. Hong, Z. Wang, X. Wei, J. Shi, and C. Li, “Antibodies against polyethylene glycol in human blood: A literature review,” J. Pharmacol. Toxicol. Methods, vol. 102, p. 106678, 2020, doi: 10.1016/j.vascn.2020.106678.

[36] T. M. Panknin, C. L. Howe, M. Hauer, B. Bucchireddigari, A. M. Rossi, and J. L. Funk, “Curcumin Supplementation and Human Disease: A Scoping Review of Clinical Trials,” Int. J. Mol. Sci., vol. 24, no. 5, p. 4476, Feb. 2023, doi: 10.3390/ijms24054476.

[37] A. B. Kunnumakkara et al., “Role of Turmeric and Curcumin in Prevention and Treatment of Chronic Diseases: Lessons Learned from Clinical Trials,” ACS Pharmacol. Transl. Sci., vol. 6, no. 4, pp. 447–518, Apr. 2023, doi: 10.1021/acsptsci.2c00012.

[38] N. Naghsh et al., “Profiling Inflammatory Biomarkers following Curcumin Supplementation: An Umbrella Meta-Analysis of Randomized Clinical Trials,” Evid.-Based Complement. Altern. Med. ECAM, vol. 2023, p. 4875636, 2023, doi: 10.1155/2023/4875636.

[39] K. Mansouri et al., “Clinical effects of curcumin in enhancing cancer therapy: A systematic review,” BMC Cancer, vol. 20, no. 1, p. 791, Aug. 2020, doi: 10.1186/s12885-020-07256-8.

[40] S. Kang, M. Kim, H. Kim, and J. Hong, “Enhancement of Solubility, Stability, Cellular Uptake, and Bioactivity of Curcumin by Polyvinyl Alcohol,” Int. J. Mol. Sci., vol. 25, no. 11, p. 6278, Jun. 2024, doi: 10.3390/ijms25116278.

[41] Y. Djoumbou Feunang et al., “ClassyFire: automated chemical classification with a comprehensive, computable taxonomy,” J. Cheminformatics, vol. 8, p. 61, 2016, doi: 10.1186/s13321-016-0174-y.

[42] Y. K. Kim, J. H. Hwang, and H. T. Lee, “Differential susceptibility to lipopolysaccharide affects the activation of toll-like-receptor 4 signaling in THP-1 cells and PMA-differentiated THP-1 cells,” Innate Immun., vol. 28, no. 3–4, pp. 122–129, Apr. 2022, doi: 10.1177/17534259221100170.

[43] A. H. Sneharani, “Curcumin-sunflower protein nanoparticles-A potential antiinflammatory agent,” J. Food Biochem., vol. 43, no. 8, p. e12909, Aug. 2019, doi: 10.1111/jfbc.12909.

[44] A. C. V. de Guzman, M. A. Razzak, J. H. Cho, J. Y. Kim, and S. S. Choi, “Curcumin-Loaded Human Serum Albumin Nanoparticles Prevent Parkinson’s Disease-like Symptoms in C. elegans,” Nanomaterials, vol. 12, no. 5, p. 758, Feb. 2022, doi: 10.3390/nano12050758.

[45] L. Fan, Y. Lu, X.-K. Ouyang, and J. Ling, “Development and characterization of soybean protein isolate and fucoidan nanoparticles for curcumin encapsulation,” Int. J. Biol. Macromol., vol. 169, pp. 194–205, Feb. 2021, doi: 10.1016/j.ijbiomac.2020.12.086.

[46] A. B. Sravani, E. M. Mathew, V. Ghate, and S. A. Lewis, “A Sensitive Spectrofluorimetric Method for Curcumin Analysis,” J. Fluoresc., vol. 32, no. 4, pp. 1517–1527, 2022, doi: 10.1007/s10895-022-02947-w.

[47] “Curcumin-Loaded Lipid Nanocarriers: A Targeted Approach for Combating Oxidative Stress in Skin Applications - PubMed.” Accessed: Feb. 02, 2026. [Online]. Available: https://pubmed-ncbi-nlm-nih-gov.ezproxy.library.dal.ca/40006512/

[48] P.-C. Li et al., “Gelatin scaffold with multifunctional curcumin-loaded lipid-PLGA hybrid microparticles for regenerating corneal endothelium,” Mater. Sci. Eng. C Mater. Biol. Appl., vol. 120, p. 111753, Jan. 2021, doi: 10.1016/j.msec.2020.111753.

[49] X.-H. Do et al., “Differential Cytotoxicity of Curcumin-Loaded Micelles on Human Tumor and Stromal Cells,” Int. J. Mol. Sci., vol. 23, no. 20, p. 12362, Oct. 2022, doi: 10.3390/ijms232012362.

[50] P. Gupta, C. T. Jordan, M. I. Mitov, D. A. Butterfield, J. Z. Hilt, and T. D. Dziubla, “Controlled curcumin release via conjugation into PBAE nanogels enhances mitochondrial protection against oxidative stress,” Int. J. Pharm., vol. 511, no. 2, pp. 1012–1021, Sep. 2016, doi: 10.1016/j.ijpharm.2016.07.071.

[51] H. Li et al., “Curcumin protects against cytotoxic and inflammatory effects of quartz particles but causes oxidative DNA damage in a rat lung epithelial cell line,” Toxicol. Appl. Pharmacol., vol. 227, no. 1, pp. 115–124, Feb. 2008, doi: 10.1016/j.taap.2007.10.002.

[52] S. Sunoqrot, B. Orainee, D. A. Alqudah, F. Daoud, and W. Alshaer, “Curcumin-tannic acid-poloxamer nanoassemblies enhance curcumin’s uptake and bioactivity against cancer cells in vitro,” Int. J. Pharm., vol. 610, p. 121255, Dec. 2021, doi: 10.1016/j.ijpharm.2021.121255.

[53] P. S. Jahagirdar, P. K. Gupta, S. P. Kulkarni, and P. V. Devarajan, “Polymeric curcumin nanoparticles by a facile in situ method for macrophage targeted delivery,” Bioeng. Transl. Med., vol. 4, no. 1, pp. 141–151, Jan. 2019, doi: 10.1002/btm2.10112.

[54] A. Safdar et al., “The Shell Thickness of Polydopamine Nanocapsules Influences Protein Adsorption and Cellular Uptake,” Biomacromolecules, Jan. 2026, doi: 10.1021/acs.biomac.5c02050.

[55] W. S. Sarawi, A. M. Alhusaini, H. K. Alghibiwi, J. S. Alsaab, and I. H. Hasan, “Roles of Nrf2/HO-1 and ICAM-1 in the Protective Effect of Nano-Curcumin against Copper-Induced Lung Injury,” Int. J. Mol. Sci., vol. 24, no. 18, p. 13975, Sep. 2023, doi: 10.3390/ijms241813975.

[56] W.-H. Lee, C.-Y. Loo, P. M. Young, R. Rohanizadeh, and D. Traini, “Curcumin Nanoparticles Attenuate Production of Pro-inflammatory Markers in Lipopolysaccharide-Induced Macrophages,” Pharm. Res., vol. 33, no. 2, pp. 315–327, Feb. 2016, doi: 10.1007/s11095-015-1789-9.

[57] S. K. Sandur et al., “Curcumin, demethoxycurcumin, bisdemethoxycurcumin, tetrahydrocurcumin and turmerones differentially regulate anti-inflammatory and anti-proliferative responses through a ROS-independent mechanism,” Carcinogenesis, vol. 28, no. 8, pp. 1765–1773, Aug. 2007, doi: 10.1093/carcin/bgm123.

[58] R. L. Edwards et al., “Mechanistic Differences in the Inhibition of NF-κB by Turmeric and Its Curcuminoid Constituents,” J. Agric. Food Chem., vol. 68, no. 22, pp. 6154–6160, Jun. 2020, doi: 10.1021/acs.jafc.0c02607.

[59] G. S. Jeong et al., “Comparative effects of curcuminoids on endothelial heme oxygenase-1 expression: ortho-methoxy groups are essential to enhance heme oxygenase activity and protection,” Exp. Mol. Med., vol. 38, no. 4, pp. 393–400, Aug. 2006, doi: 10.1038/emm.2006.46.

