## Supplemental Figures for "Turmeric Phyto-NanoParticle (TPNP) enhances cellular bioavailability and anti-inflammatory effect of curcuminoids in human monocytes / macrophages"

Figure S1

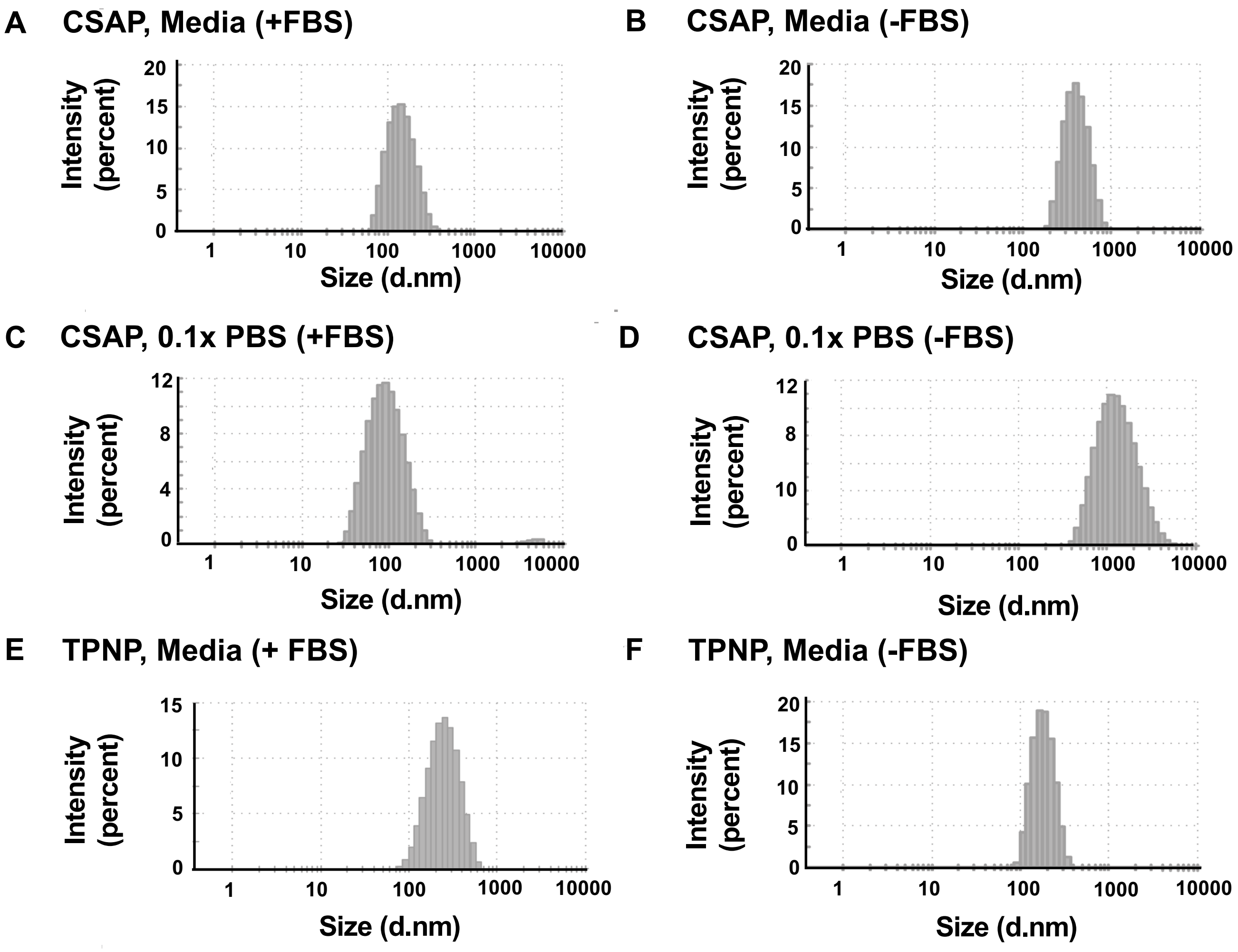
A
CSAP, Media (+FBS)
B
CSAP, Media (-FBS)
C
CSAP, 0.1x PBS (+FBS)
D
CSAP, 0.1x PBS (-FBS)
E
TPNP, Media (+ FBS)
F
TPNP, Media (-FBS)
Intensity (percent)
Size (d.nm)
1
10
100
1000
10000
0
5
10
15
20
0
4
8
12

Figure S2

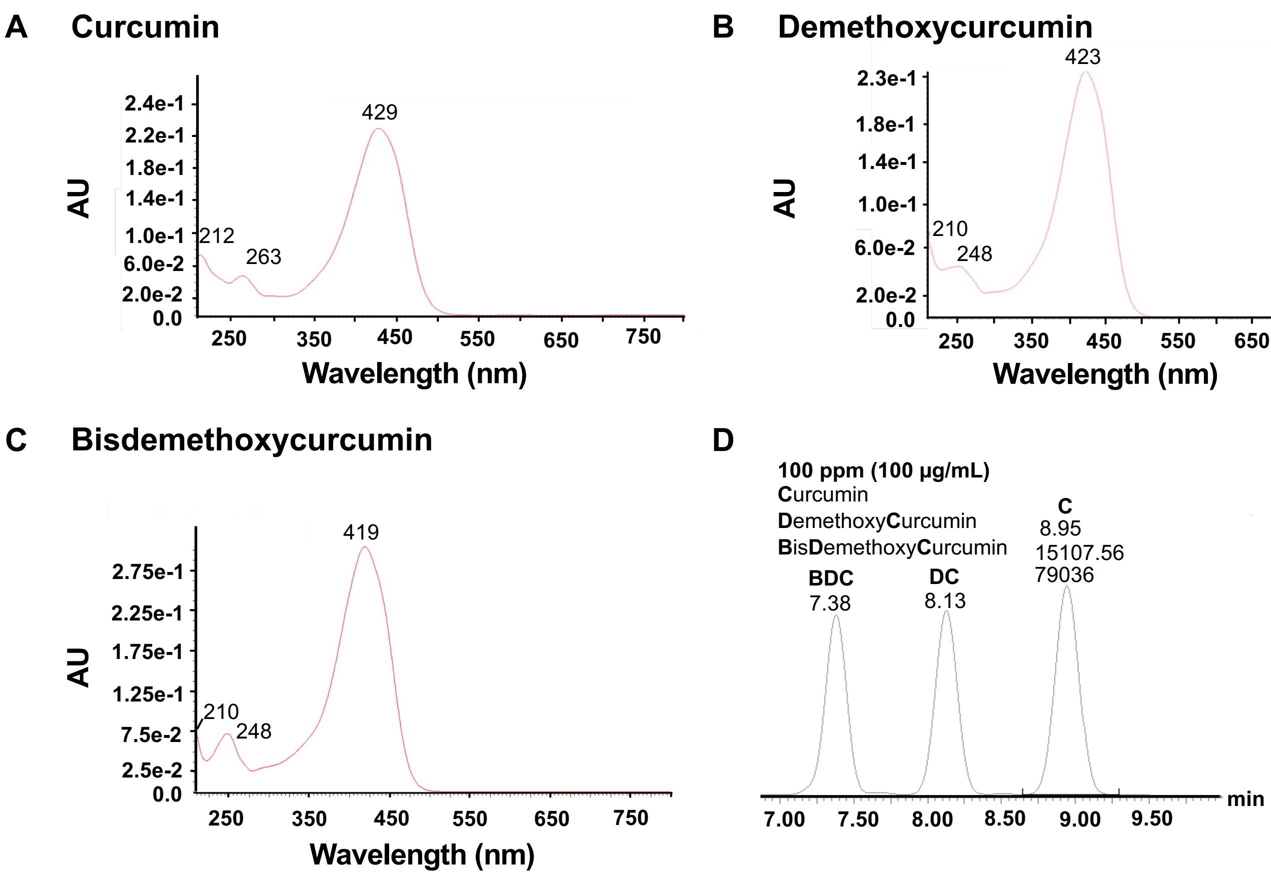
A
Curcumin
AU
2.4e-1
2.2e-1
1.8e-1
1.4e-1
1.0e-1
6.0e-2
2.0e-2
0.0
212
263
429
250
350
450
550
650
750
Wavelength (nm)
B
Demethoxycurcumin
AU
2.3e-1
1.8e-1
1.4e-1
1.0e-1
6.0e-2
2.0e-2
0.0
210
248
423
250
350
450
550
650
750
Wavelength (nm)
C
Bisdemethoxycurcumin
AU
2.75e-1
2.25e-1
1.75e-1
1.25e-1
7.5e-2
2.5e-2
0.0
210
248
419
250
350
450
550
650
750
Wavelength (nm)
D
100 ppm (100 μg/mL)
Curcumin
DemethoxyCurcumin
BisDemethoxyCurcumin
BDC
7.38
DC
8.13
C
8.95
15107.56
79036
7.00
7.50
8.00
8.50
9.00
9.50
min

Figure S3

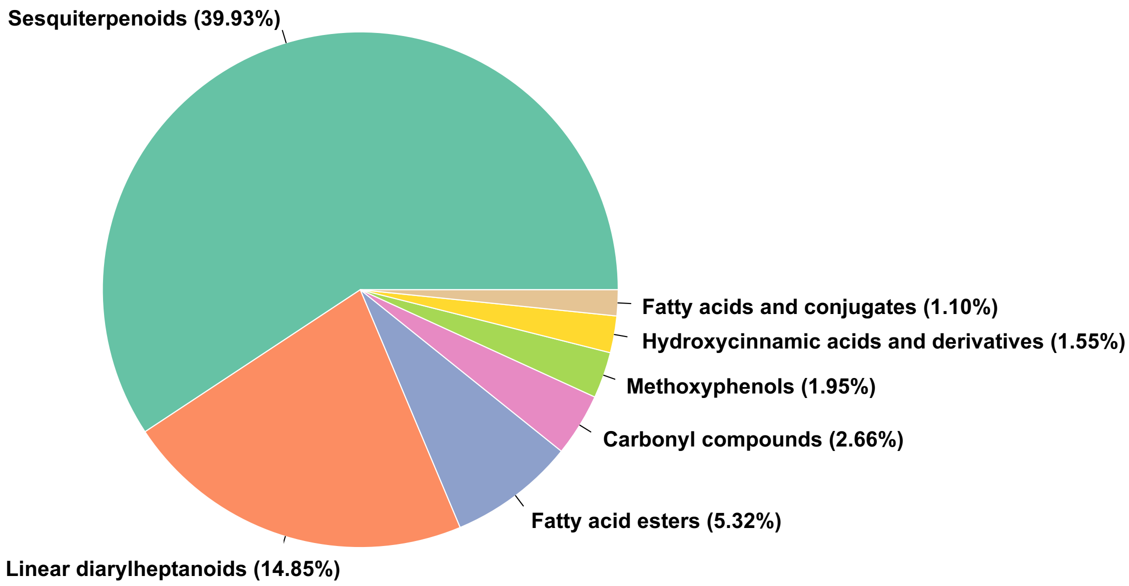
Sesquiterpenoids (39.93%)
Fatty acids and conjugates (1.10%)
Hydroxycinnamic acids and derivatives (1.55%)
Methoxyphenols (1.95%)
Carbonyl compounds (2.66%)
Fatty acid esters (5.32%)
Linear diarylheptanoids (14.85%)

Figure S4

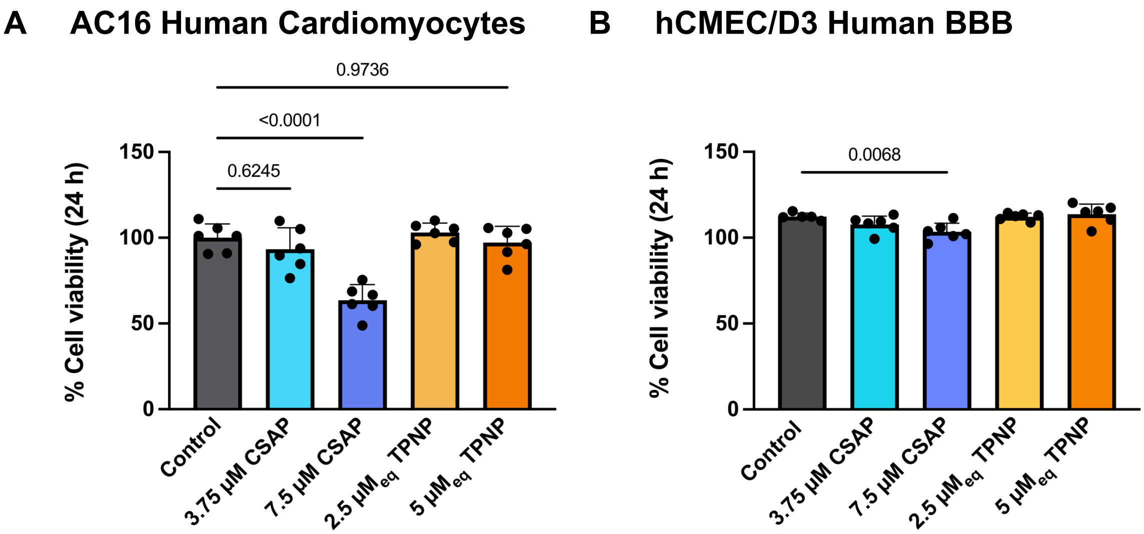
A
AC16 Human Cardiomyocytes
0.9736
<0.0001
0.6245
% Cell viability (24 h)
150
100
50
0
Control
3.75 µM CSAP
7.5 µM CSAP
2.5 µMeq TPNP
5 µMeq TPNP
B
hCMEC/D3 Human BBB
0.0068
% Cell viability (24 h)
150
100
50
0
Control
3.75 µM CSAP
7.5 µM CSAP
2.5 µMeq TPNP
5 µMeq TPNP

Figure S5

**A** **Monocytes**

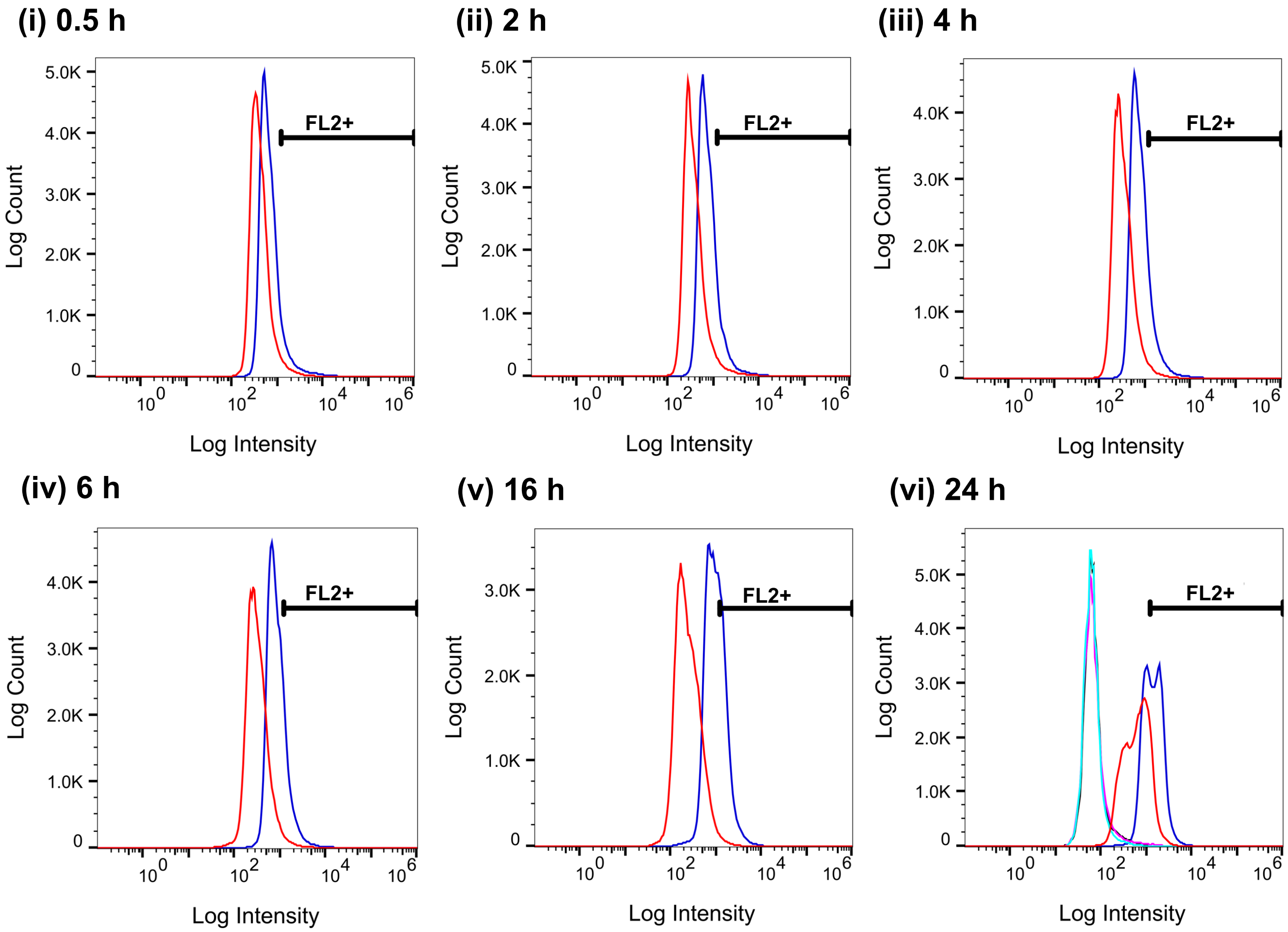


FACS Legend: 7.5 µM CSAP; 5 $\mu M_{eq}$ TPNP; Cells only; TPNP veh; EtOH veh

Table S1

| Macrophages | | | | |
|---|---|---|---|---|
| **Sample** | **Time (h)** | **% Total** | **Median** | **Gate (%)** |
| **Media only** | 24 | 100 | 87.2 | 0.26 |
| **EtOH veh** | 24 | 100 | 92.1 | 0.41 |
| **TPNP veh** | 24 | 100 | 96.9 | 0.35 |
| **7.5 µM CSAP** | 0.5 | 100 | 277 | 8.05 |
| | 1 | 100 | 370 | 20.2 |
| | 2 | 100 | 423 | 28.8 |
| | 4 | 100 | 372 | 19.7 |
| | 6 | 100 | 349 | 17.4 |
| | 16 | 100 | 236 | 5.58 |
| | 24 | 100 | 310 | 16.9 |
| **5 $\mu M_{eq}$ TPNP** | 0.5 | 100 | 527 | 47.5 |
| | 1 | 100 | 636 | 65.6 |
| | 2 | 100 | 763 | 79.6 |
| | 4 | 100 | 872 | 87.7 |
| | 6 | 100 | 1028 | 92.4 |
| | 16 | 100 | 1271 | 95.3 |
| | 24 | 100 | 1112 | 89.8 |

Table S2

| Monocytes | | | | |
|---|---|---|---|---|
| **Sample** | **Time (h)** | **% Total** | **Median** | **Gate (%)** |
| **Cells only** | 24 | 100 | 64.9 | 0.27 |
| **EtOH veh** | 24 | 100 | 63.9 | 0.70 |
| **TPNP veh** | 24 | 100 | 62.9 | 0.15 |
| **7.5 µM CSAP** | 0.5 | 100 | 389 | 4.08 |
| | 1 | 100 | 380 | 3.21 |
| | 2 | 100 | 345 | 4.28 |
| | 4 | 100 | 313 | 2.85 |
| | 6 | 100 | 303 | 2.51 |
| | 16 | 100 | 228 | 1.42 |
| | 24 | 100 | 659 | 15.4 |
| **5 $\mu M_{eq}$ TPNP** | 0.5 | 100 | 622 | 11.2 |
| | 1 | 100 | 593 | 8.10 |
| | 2 | 100 | 710 | 15.0 |
| | 4 | 100 | 739 | 16.9 |
| | 6 | 100 | 803 | 20.2 |
| | 16 | 100 | 962 | 33.6 |
| | 24 | 100 | 1397 | 59.4 |
